# Linking signal detection theory and encoding models to reveal independent neural representations from neuroimaging data

**DOI:** 10.1101/254995

**Authors:** Fabian A. Soto, Lauren E. Vucovich, F. G. Ashby

## Abstract

Many research questions in visual perception involve determining whether stimulus properties are represented and processed independently. In visual neuroscience, there is great interest in determining whether important object dimensions are represented independently in the brain. For example, theories of face recognition have proposed either completely or partially independent processing of identity and emotional expression. Unfortunately, most previous research has only vaguely defined what is meant by “independence,” which hinders its precise quantification and testing. This article develops a new quantitative framework that links signal detection theory from psychophysics and encoding models from computational neuroscience, focusing on a special form of independence defined in the psychophysics literature: perceptual separability. The new theory allowed us, for the first time, to precisely define separability of neural representations and to theoretically link behavioral and brain measures of separability. The framework formally specifies the relation between these different levels of perceptual and brain representation, providing the tools for a truly integrative research approach. In particular, the theory identifies exactly what valid inferences can be made about independent encoding of stimulus dimensions from the results of multivariate analyses of neuroimaging data and psychophysical studies. In addition, commonly used operational tests of independence are re-interpreted within this new theoretical framework, providing insights on their correct use and interpretation. Finally, we apply this new framework to the study of separability of brain representations of face identity and emotional expression (neutral/sad) in a human fMRI study with male and female participants.

**Author Summary:** A common question in vision research is whether certain stimulus properties, like face identity and expression, are represented and processed independently. We develop a theoretical framework that allowed us, for the first time, to link behavioral and brain measures of independence. Unlike previous approaches, our framework formally specifies the relation between these different levels of perceptual and brain representation, providing the tools for a truly integrative research approach in the study of independence. This allows to identify what kind of inferences can be made about brain representations from multivariate analyses of neuroimaging data or psychophysical studies. We apply this framework to the study of independent processing of face identity and expression.

## Introduction

A common goal in perceptual science is to determine whether some stimulus dimensions or components are processed and represented independently from other types of information. In visual neuroscience, much research has focused on determining whether there is independent processing of object and spatial visual information [1], of object shape and viewpoint [2], of different face dimensions [3, 4], etc. A common approach is to use operational definitions of independence, which are linked to rather vague conceptual definitions. This approach has the disadvantage that different researchers use different operational definitions for independence, often leading to contradictory conclusions. For example, in the study of whether face identity and emotional expression are processed independently, evidence for both independence and interactivity has been found across a variety of operational tests. Evidence for independence was found by most lesion studies [5], by lack of correlation between fMRI patterns related to identity and expression [6], by single neuron invariance [7], by selective fMRI adaptation release in fusiform face area (FFA) and middle superior temporal sulcus (STS) [8], and by selective fMRI decoding of identity from anterior FFA and medial temporal gyrus [9, 10], and of expression from STS [10]. Evidence for a lack of independence has been provided by overlapping fMRI activation during filtering tasks [11], by non-selective fMRI adaptation release in posterior STS [8] and in FFA-when adaptation is calculated based on perception [12]-, and by non-selective fMRI decoding from right FFA [9].

Because the different operational definitions are not linked to one another through a theoretical framework, the interpretation of such contradictory results is very difficult and necessarily post-hoc. Even more difficult is to link the neurobiological results to the psychophysics literature on independence of face dimensions, which itself is plagued by similar issues [for a review, see 13].

General recognition theory (GRT) [14, 15] is a multidimensional extension of signal detection theory that has solved such problems in psychophysics, by providing a unified theoretical framework in which notions of independence can be defined and linked to operational tests. Hundreds of studies have applied GRT to a wide variety of phenomena, including face perception [16, 17], recognition and source memory [18, 19], source monitoring [20], object recognition [21, 22], perception/action interactions [23], speech perception [24], haptic perception [25], the perception of sexual interest [26], and many others.

Here we present an extension of GRT to the study of independence of brain representations, by relating it to encoding models and decoding methods from computational neuroscience [27, 28]. Past neuroimaging studies have been limited to choosing between decoding methods, which try to determine what stimulus information is processed in a brain region while ignoring the form of the underlying representation, and encoding models, which assume a specific representation and compare its predictions against data. [29]. We propose the concept of encoding separability as a fundamental way in which brain representations of stimulus properties can be considered independent, and we identify the specific conditions in which a decoding analysis of neuroimaging data or a psychophysical study allow inferences to be made about encoding separability. In doing so, we show that decoding methods (and under some assumptions, psychophysics) can be useful to make valid inferences about encoding. We also re-interpret previously-proposed tests of independence within our new theoretical framework, and provide guides on their correct use. Finally, we apply this new framework to the study of separability of brain representations of face identity and expression.

## Results

### Extending General Recognition Theory to the Study of Brain Representations

GRT is a multivariate extension of signal detection theory to cases in which stimuli vary on more than one dimension [14, 15]. As in signal detection theory, the theory assumes that different presentations of the same stimulus produce slightly different perceptual representations. For example, as shown in Figure 1, repeated presentations of a face identity produce a variety of values on the “identity” dimension (orange and red dots), which follow a probability distribution (red and orange curves). According to GRT, there are many ways in which processing of a dimension of interest, or target dimension, can be influenced by variations in a second, irrelevant dimension. GRT formally defines such dimensional interactions and links them to operational tests of independence. This allows researchers to determine whether a test can dissociate between different forms of independence, and to create new tests specifically designed to target a specific form of independence.

**Figure 1:**
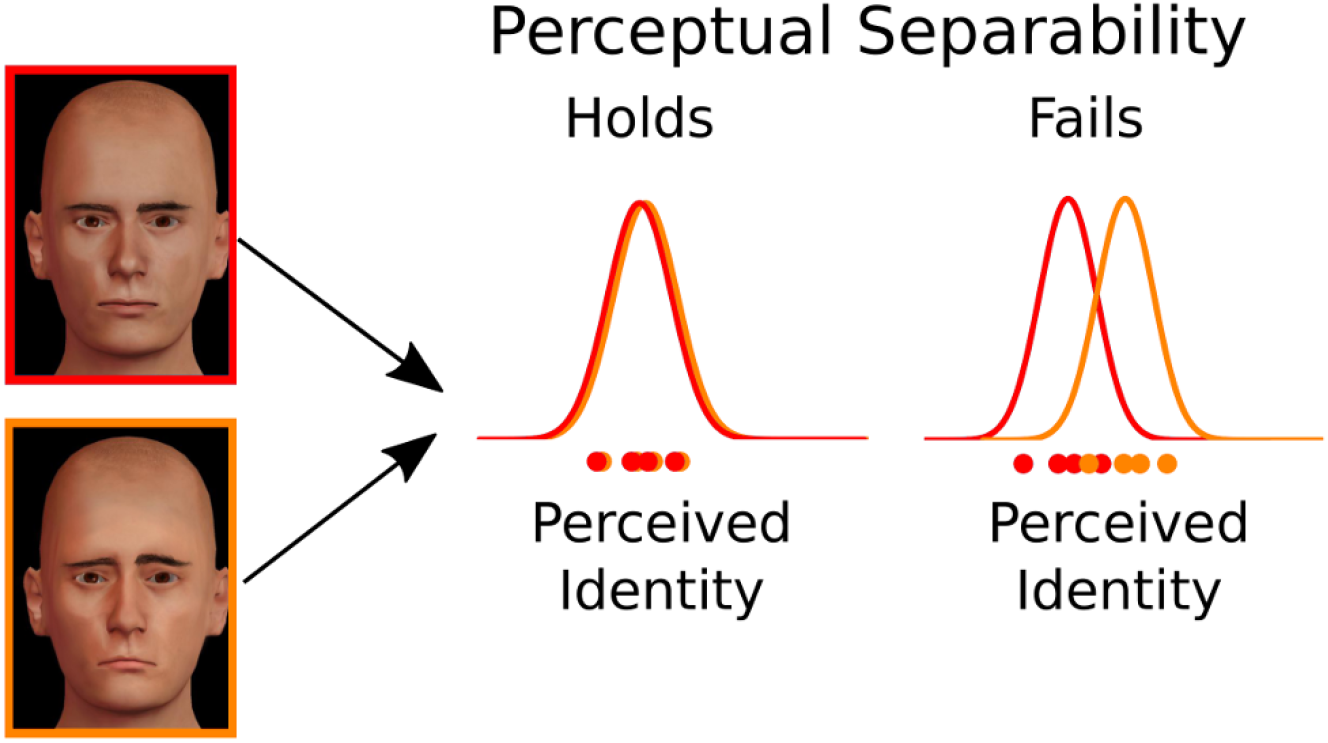
Stimulus representation and definition of perceptual separability in GRT. The representation of a given identity changes randomly from trial to trial (dots at the bottom) according to some perceptual distribution (bell-shaped distributions at the top). Perceptual separability of identity from emotional expression (neutral vs. sad) holds if the perceptual distribution for identity does not change with emotional expression (left), and it fails if the perceptual distribution for identity does change with emotional expression (right).

Here we will consider the special case in which stimuli vary along two stimulus dimensions (or more generally, components or properties), represented by *A* and *B*. However, the theory can easily be extended to a larger number of dimensions. Specific values of dimension A used in an experiment are indexed by *i* = 1, 2,…*L_A_*, and the specific values of dimension *B* are indexed by *j* = 1, 2, … *L_B_*. A stimulus in the experiment is represented by a combination of these dimension levels, *A_i_B_j_*. This stimulus produces a random perceptual effect in a two-dimensional perceptual space [*x, y*], where *x* represents the perceptual effect of property *A* and *y* the perceptual effect of property *B*. The random vector [*x,y*] can be described through a two-dimensional joint probability density *p*(*x, y|A_i_B_j_*), with *p*(*x|A_i_B_j_*) and *p*(*y|A_i_B_j_*) representing the marginal densities of the perceptual effects associated with components *A* and *B*, respectively (the distributions shown in Figure 1 are examples of such marginal densities).

#### Perceptual separability and perceptual independence

A particularly important form of independence defined in GRT is *perceptual separability*, which holds when the perception of the target dimension is not affected by variations in the irrelevant dimension. In Figure 1, an identity is presented with a neutral expression (in orange) or with a sad expression (in red). When perceptual separability holds, the orange and red perceptual distributions overlap, and the face is just as easy to identify in both cases. When perceptual separability fails, the orange and red perceptual distributions do not overlap, and the face is easier to identify when the expression is sad (there is more evidence for the identity in this case).

Here we focus on perceptual separability because it is considered a particularly important form of independence, for two reasons. First, because many questions in perceptual neuroscience can be understood as questions about separability of object dimensions. For example, the question of whether object representations are invariant across changes in identity-preserving variables like rotation and translation is equivalent to the question of whether object representations are perceptually separable from such variables [30]. In face perception, configural or holistic face perception has been defined as non-separable processing of different face features [31, 32], and the question of whether or not different face dimensions are processed independently is usually investigated using tests of perceptual separability [33, 34, 35, 13]. The second reason for the importance of perceptual separability is that higher-level cognitive mechanisms seem to be applied differently when stimuli differ along separable dimensions rather than along non-separable dimensions. For example, selective attention is deployed more easily to separable dimensions than to non-separable dimensions [36, 37], sources of predictive and causal knowledge may be combined differently if they differ along separable versus non-separable dimensions [38, 39], and the performance cost of storing objects in visual working memory is different depending on whether such objects differ from one another in separable versus non-separable dimensions [40].

Formally, perceptual separability of dimension *A* from dimension *B* occurs when the perceptual effect of stimuli on dimension *A* does not change with the value of the stimulus on dimension *B* [14]-that is, if and only if, for all values of *x* and *i*:

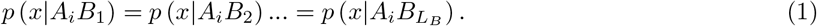

Perceptual separability of dimension *B* from dimension *A* is defined analogously.

Another form of independence defined within GRT is *perceptual independence*. Perceptual independence of components A and B holds in stimulus *A_i_B_j_* if and only if the perceptual effects of A and B are statistically independent; that is, if and only if:

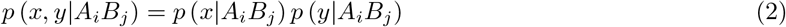

Unlike perceptual separability, which is a form of independence involving the representation of multiple stimuli, perceptual independence refers to dimensional interactions in the representation of a single stimulus.

#### Neural encoding and encoding separability

Extending GRT to the study of neural representation requires linking it to our current understanding on how dimensions are represented by neuronal populations. In the computational neuroscience literature, an encoding model is a formal representation of the relation between sensory stimuli and the response of a single neuron or a group of neurons [41, 28, 42]. In the case of stimulus dimensions, an encoding model represents how changes in a dimension of interest are related to changes in neural responses. Encoding models have been applied to describe neural responses at a variety of scales, from single neurons to the average activity of thousands of neurons [43, 41, 28, 29]. To discuss these models in their more general form, it is convenient to introduce the abstract concept of a *channel*, which can be used as a placeholder for a single neuron, a population of neurons with similar properties, or as an abstract construct to model the behavior of a human observer. A channel is essentially a detector, sensitive to a particular stimulation pattern. It responds maximally to that target pattern and progressively less to other patterns as they become different from the target. In other words, the most important property of a channel is that it has tuning. The tuning of a channel can be modeled in many ways, but perhaps the simplest is to choose a physical dimension of interest and model the channel’s response as a function of the value of a stimulus on that dimension. For example, if we are interested in dimension *A* (equivalent definitions can be given for *B*), then the response *r_c_* of the channel *c* to a stimulus *A_i_B_j_* is determined by a tuning function:

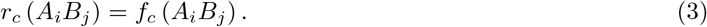

Common choices for *f_c_*(*A_i_B_j_*) in the literature are bell-shaped and sigmoidal functions [28]. The channel response on a given trial may also be influenced by stochastic internal noise, which can be assumed to be additive (independent of the channel’s response) or multiplicative (scaling with the channel’s response). Common choices for the distribution of this noise in the literature are Gaussian and Poisson [28, 42]. Because the noise is a random variable, the response of the channel *r_c_* itself becomes a random variable that follows a probability distribution:

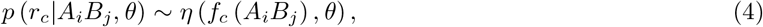

where ~ means “distributed as”, and *η*() is just a placeholder that stands for any probability distribution (e.g., Gaussian) that depends on the channel’s tuning function and on a set of parameters *θ* describing noise.

Researchers agree that encoding of a stimulus dimension requires a model with multiple channels, or multichannel model. For example, the “standard model” of dimension encoding in the computational neuroscience literature is such a multichannel model (implementing a “population code” [41, 28]), and most applications in the neuroscience and psychophysics literature use at least two channels to describe encoding of stimulus dimensions [44]. Figure 2 shows encoding of a stimulus dimension with four channels, each with its own tuning model represented by a curve of different color. The tuning model is a formalization of how the channel responds to different stimulus values: each channel responds maximally to its preferred dimensional value and less to other values. The figure shows the response of a multi-channel model to a stimulus with a value of 3 on the target dimension. A channel’s noise model describes the stochasticity in the channel’s responses through a probability distribution. In Figure 2, the average response of each channel is perturbed by random additive noise, represented by the dice. The final channel output is equal to the average response (from the tuning model) plus noise (randomly drawn from the noise model).

**Figure 2:**
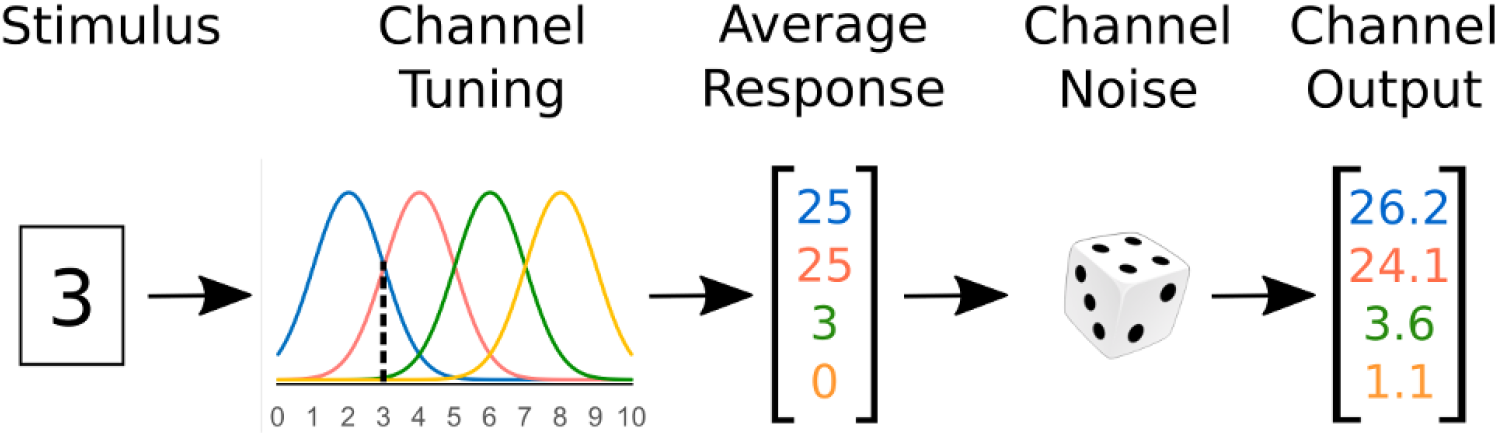
Schematic representation of multiple channels encoding a stimulus with a value of “3” in a target dimension. If a stimulus with value “3” is presented, each channel gives an average response equivalent to the height of the tuning function at that stimulus value (i.e., the height at the dotted line). The vector of average responses is perturbed by random noise, producing the final channel output.

A multichannel model encodes information about dimension A through the combined response of *N* channels [41, 28, 44], indexed by *c* =1, 2,…*N*. On each trial, the model produces a (column) random vector of channel responses **r** = [*r*_1_,*r*_2_,… *r_N_*]^**T**^, where ^**T**^ denotes matrix transpose. Note that **r** depends on the stimulus *A_i_B_j_* according to a set of tuning functions:

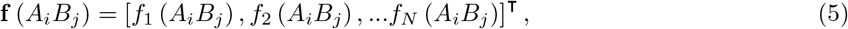

The probability distribution of **r** also depends on noise parameters *θ*:

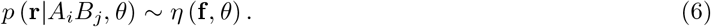

As indicated earlier, additive Gaussian noise is a common choice for the channel noise model. In that case, the multichannel encoding model is described by a multivariate Gaussian distribution:

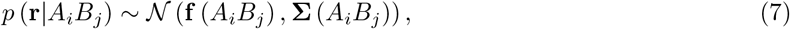

where Σ(*A_i_B_j_*) is an *N × N* covariance matrix describing channel noise. In most applications, noise is also assumed to be independently distributed across channels, and all non-diagonal cells in Σ(*A_i_B_j_*) are zero.

This discussion suggests that a new form of separability can be defined for the neural representation of a dimension: *encoding separability*. When a target dimension is encoded in the exact same way across variations of an irrelevant dimension, we say that the former shows encoding separability from the latter. For encoding separability to hold, both the tuning and noise models of all channels must be equivalent across changes in the irrelevant dimension, which is equivalent to having a single encoding model representing the target dimension, independently of the value of the irrelevant dimension.

Formally, encoding separability of dimension *A* from dimension *B* holds when encoding of the value of *A* does not change with the stimulus’ value on *B*. That is, if and only if, for all values of **r** and *i*:

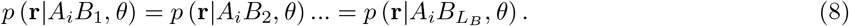

Encoding separability of dimension *B* from dimension *A* is defined analogously.

Violations of encoding separability can happen for two reasons. The first possibility is that one or more of the tuning functions in **f** change with the value of *B. Tuning separability* of dimension *A* from dimension *B* holds when all tuning functions that encode dimension *A* depend only on the value of *A*-that is, if and only if, for all channels *c* and stimuli *A_i_B_j_*:

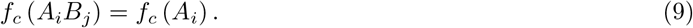

Tuning separability of dimension *B* from dimension *A* is defined analogously. Because *p*(**r**|*A_i_B_j_, θ*) depends on f (see Equation 6), violations of tuning separability produce violations of encoding separability.

The second reason for a violation of encoding separability is that the noise for one or more channels is distributed differently for different levels of *B*.

Because the Gaussian encoding model described in Equation 7 is completely characterized by the mean vector **f**(*A_i_B_j_*) and the covariance matrix Σ(*A_i_B_j_*), encoding separability of *A* from *B* holds if the following two conditions are true for all stimuli *A_i_B_j_*:

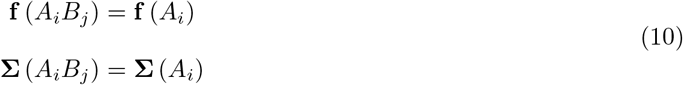

Encoding separability is a concept describing the way in which stimulus information is represented by single neurons or populations of neurons with similar characteristics. Thus, it can be directly tested only when we have access to direct measurements of **r** (e.g., firing rates from single cell recordings or a measure of the activity of a homogeneous neural population), and a sample size large enough to precisely estimate *p*(**r**|*A_i_B_j_, θ*) for all values of *i* and *j*. However, in most cases we do not have access to such direct measurements, but to indirect measures of neural activity contaminated with measurement error, as is the case in fMRI and EEG experiments. Measures of activity in neuroimaging studies result from an unknown, perhaps non-linear transformation of **r**. The encoding distributions *p*(**r**|*A_i_B_j_, θ*) cannot be estimated from such indirect measures, but we will show that there are ways to make valid inferences about encoding separability from indirect tests. For that, we must first introduce the concepts of *decoding* and *decoding separability*.

#### Decoding separability

The term neural decoding refers both to a series of methods used by researchers to extract information about a stimulus from neural data [29, 45] and to the mechanisms used by readout neurons to extract similar information, which is later used for decision making and other cognitive processes [28, 46]. If dimension *A* is encoded by *N* channels, according to the scheme summarized in Equation 6 and depicted in Figure 2, then the decoded estimate of a dimensional value *Â*, will be some function of the channel responses:

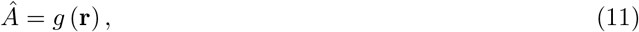

where *g*() is a function from 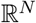 to 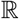 (i.e., from the multidimensional space of the channel responses to the unidimensional space of the decoded dimension). Because **r** is a random vector (see Equation 6), the decoded value *Â* is a random value that follows a probability distribution *p*(*Â|A_i_B_j_, θ*). In many cases, knowledge about the encoding distribution from Equation 6 and the decoder from Equation 11 allows one to derive an expression for *p*(*Â|A_i_B_j_, θ*). Note also that *Â* is a continuous variable, despite the fact that it is estimated as a response to stimuli with discrete levels in the stimulus dimension *A*.

There are many possible decoding schemes, but the most popular among researchers [46, 47], due to their simplicity and neurobiological plausibility, are simple linear decoders

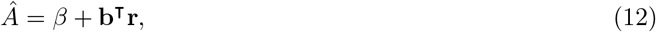

where *β*x is a scalar and **b** is a (column) vector of weights.

With a Gaussian encoding model like the one described by Equation 7, the distribution of linearly-decoded estimates of values on dimension *A* is:

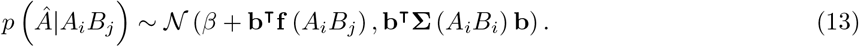

When channel noise is independent, the variance of the decoded variable *Â* is simply 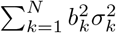, where 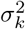 represents the *N* diagonal elements of Σ(*A_i_B_j_*).

We define *decoding separability* as the situation in which the distribution of decoded values on the target dimension is invariant across changes in the stimulus on a second, irrelevant dimension. That is, decoding separability of dimension *A* from dimension *B* holds when the distribution of decoded values of *A* does not change with the value of *B* in the stimulus-that is, if and only if, for all values of *Â* and *i*:

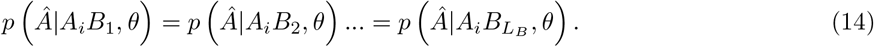

Decoding separability of dimension *B* from dimension *A* is defined analogously.

#### Relation between encoding separability and decoding separability

Decoding separability is easy to check by directly decoding dimensional values from a neuronal population. Moreover, if the same decoding scheme is used for all values of the irrelevant dimension, then the relations between encoding separability and decoding separability shown in Figure 3 hold, as we show in this section.

**Figure 3:**
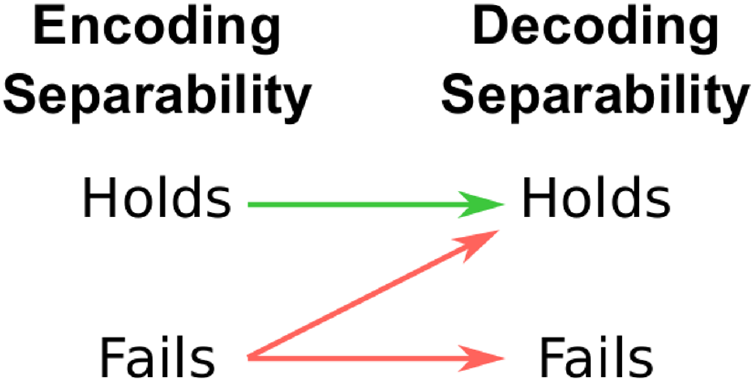
Summary of the relation between encoding separability and decoding separability, according to our extension to GRT. Arrows should be interpreted as conditional statements of the form “if X, then Y”. These relations mean that a failure of encoding separability is a valid inference from the observation of a failure of decoding separability. However, the presence of encoding separability cannot be validly inferred from an observation of decoding separability.

##### If encoding separability holds, then decoding separability must also hold

This proposition is represented by the green arrow in Figure 3. When encoding separability holds (see Equation 8), *p*(**r**|*A_i_B_j_,θ*) = *p*(**r**|*A_i_,θ*) for all values of **r** and *j*. Because we have assumed that decoding depends only on the value of **r** (Equation 11), the function *g*() is also independent of the value of *B_j_*. Thus, regardless of the shape of *p*(**r**|*A_i_, θ*) and *g*(), the distribution of the decoded variable *Â* is independent of the value of *B_j_*, and decoding separability (Equation 14) holds. In other words, for all values of Bj the same decoding transformation *g*() is applied to the same encoding distribution *p*(**r**|*A_i_,θ*), resulting in the same decoding distribution *p*(*Â|A_i_, θ*).

##### If encoding separability fails, then decoding separability may fail or hold

This is represented by the red arrows in Figure 3. Our strategy to prove this proposition will be to disprove two universal statements through counterexamples.

We start by offering a counterexample disproving the following universal statement: *if encoding separability fails, then decoding separability must fail*. Suppose that a dimension *A* is encoded through the model with Gaussian channel noise described by Equation 7, and that we use a linear decoder to estimate *Â*, as described by Equation 12. Also suppose that there are violations of tuning separability (Equation 9) of *A* from *B* in the encoding model. Without loss of generality, suppose that those violations are differences in the tuning functions of *A*_1_*B*_1_ and *A*_1_*B*_2_:

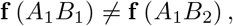

or equivalently

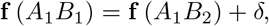

where *δ* represents a *N* × 1 vector of deviations from tuning separability, and *δ* ≠ 0.

Under the assumptions listed above, the tuning functions only affect the mean of the decoded variable (see Equation 13), so we can ignore its variance. Now suppose that in this model decoding separability holds. In that case, we have that:

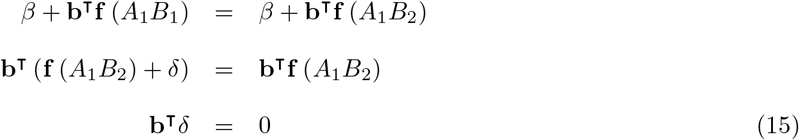

For any given *δ* ≠ **0**, there are an infinite number of **b** ≠ **0** that satisfy this equation, yielding a model in which encoding separability fails and decoding separability holds. The universal statement *if encoding separability fails, then decoding separability must fail* is false.

We now offer a counterexample to disprove the alternate universal statement: *if encoding separability fails, then decoding separability must hold*. Following the same line of reasoning as before, we get that if decoding separability fails then:

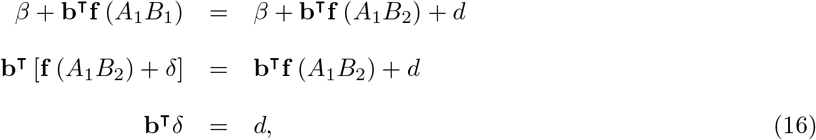

where *d* represents a scalar deviation from decoding separability in the mean of the decoding distributions. As before, for any given *δ* ≠ **0** and *d* ≠ 0, there are an infinite number of **b** ≠ 0 that satisfy this equation, yielding a model in which encoding separability fails and decoding separability fails. The universal statement *if encoding separability fails, then decoding separability must hold* is false.

In summary, if encoding separability fails, then decoding separability may hold or fail. While no universal statements can be made about decoding separability when encoding separability fails, a more specific relation may hold for particular combinations of encoding models and decoding schemes. For example, it might be that for some specific combination of an encoding model and decoding scheme, a failure of encoding separability necessarily leads to a failure of decoding separability, eliminating the diagonal arrow in Figure 3. If that was the case, it would be possible to infer encoding separability from a finding of decoding separability. We will not explore these possibilities here and instead will leave them for future work. However, note that the counterexamples offered here involve normally-distributed channel noise and a linear decoder, both of which are common choices in the literature on encoding and decoding. That is, under common assumptions and methods it is not possible to infer encoding separability from a finding of decoding separability.

One general result that must hold true is the following: if encoding separability fails and *Â* = *g*(**r**) is an injective (one-to-one) mapping, then decoding separability must fail. This is the case because when encoding separability fails (without loss of generality) *p*(**r**|*A*_1_*B*_1_,*θ*) ≠ *p*(**r**|*A*_1_*B*_2_,*θ*) for at least one **r**. If *g*() is injective, then this difference in probability at **r** will translate to a difference in probability at the corresponding transformed variable *Â*. However, because *g*() is a transformation from 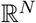 to 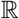, it is in most cases not injective. For example, a linear *g*() from 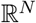 to 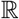 cannot be injective.

#### Inferring encoding separability from tests of decoding separability

From the results of the previous section, which are summarized in Figure 3, we can conclude that the observation of a violation of decoding separability in a particular brain region is diagnostic of a corresponding violation of encoding separability. This is because a violation of decoding separability cannot be produced when encoding separability holds. On the other hand, when decoding separability holds nothing can be concluded about encoding separability, as both encoding separability and failures of encoding separability can lead to decoding separability, depending on features of the decoder. This allows an indirect test of encoding separability, which is useful for cases where directly observing encoding separability is difficult (e.g., when indirect measures of neural activity are used, as in fMRI).

#### Perceptual separability as a form of decoding separability

The concepts of encoding and decoding separability can be linked back to GRT by assuming that perception of a dimensional value is a form of decoding. That is, the key is to assume that the perceptual representation of a stimulus dimension in GRT (the “perceived identity” in Figure 1) is the outcome of decoding a dimensional value from the activity of many channels distributed across the brain (like those shown in Figure 2). This assumption is not new and has proven useful in applications of signal detection theory in the past [44].

More specifically, assume that the perceived value of dimension *A, x*, is the result of decoding a dimensional value from the activity of many channels distributed across the brain, as in Equation 11. In other words, the perceived value *x* is a special case of the decoded variable *Â*, but estimated by readout neurons with the goal of guiding behavioral responses in a perceptual task. Denote the decoding function used by these readout neurons to obtain the perceived value *x* as *g_R_*(), so that

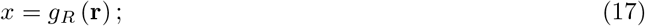

this is just a special case of the more general decoding function shown in Equation 11, but limited only to the decoding schemes that can be implemented by real neurons. We have that

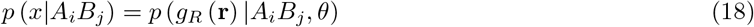

is the marginal distribution of perceptual effects along *x*. Under these assumptions, perceptual separability (Equation 1) is a form of decoding separability (Equation 14). As a consequence, from Figure 3 we know that *any failure of perceptual separability documented in the literature should be reflected in a failure of encoding separability, in brain areas providing useful information for perceptual identification of dimensional values*. The exact brain regions that provide information to solve a particular task are usually unknown, but we can assume that they encode such information in a relatively transparent (easily decodable) way. The set of potential candidates can be reduced to areas known to provide useful information for behavioral performance. Novel methods to identify such areas, which combine information about decoded values in a dimension and behavioral response times, have been recently developed [48] and seem very promising. For neuroscientists, this opens the opportunity to link new research on the separability of neural representations with decades of accumulated psychophysical research on perceptual separability [15].

In addition, as we have seen in the previous section, the common assumption of a Gaussian distribution of perceptual effects [15, 13] is met when the encoding model has additive Gaussian noise and the decoder is linear (see Equation 13), two assumptions that are common in the literature.

#### Encoding and decoding independence

Here we have focused mostly on the concept of separability. As explained earlier, this concept is particularly important because it captures features of representations, such as invariance and configurality, that are widely studied in vision science. Still, our extended GRT framework allows to define dimensional interactions that are analogous to the concept of perceptual independence from the traditional GRT, but at the level of encoding model and decoded variables. Here we define these forms of independence and very briefly discuss their relation under special assumptions about decoding. We leave a more complete treatment for future work.

Suppose that there are two different sets of channels **r**_*A*_ and **r**_*B*_, encoding stimulus components *A* and *B*, respectively. The distribution of neural responses **r**_*A*_ to stimulus *A_i_B_j_* is represented by *p*(**r**_*A*_|*A_i_B_j_, θ*), and the distribution of responses **r**_*B*_ to stimulus *A_i_B_j_* is represented by *p*(**r**_*B*_|*A_i_B_j_, θ*). Given this, *encoding independence* of components *A* and *B* holds in stimulus *A_i_B_j_* if and only if the two encoding distributions are statistically independent; that is, if and only if:

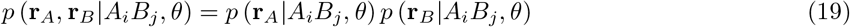

Now suppose that there are also two separate decoders, *g_A_*() to obtain estimate *Â* and *g_B_*() to obtain estimate 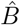. Each of these decoded values is a random variable that follows a probability distribution, represented by *p*(*Â|A_i_B_j_, θ*) and (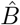. Given this, *decoding independence*|A_i_B_j_, θ) of estimates *Â* and 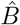 holds in stimulus *A_i_B_j_* if and only if the two decoding distributions are statistically independent; that is, if and only if:

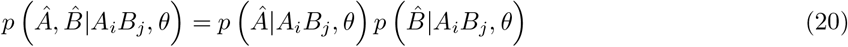

What is the relation between encoding and decoding independence? We can start by answering this question for the simple case in which the two decoding functions *g_A_* and *g_B_*, from which *Â* and 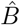 are estimated, have completely separated domains, meaning that the domain of *g_A_* does not include any of the channels in **r**_*B*_ and the domain of *g_B_* does not include any of the channels in **r**_*A*_. Under this assumption, *if encoding independence holds, then decoding independence must hold*. If **r**_*A*_ and **r**_*B*_ are independent random vectors and the decoding functions *g_A_* and *g_B_* used to obtain *Â* and 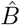 are borel-measurable functions, then *Â* and 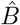 must be independent. On the other hand, *if encoding independence fails, then decoding independence may hold or fail*. We prove this statement by giving two rather trivial counterexamples to universal statements. Suppose that encoding independence fails due to a failure of pairwise independence, in which variables *r*_*A*1_ and *r*_*B*1_ are statistically dependent. Then one can choose linear *g_A_* and *g_B_* so that all or most of the variability in *Â* is due to *r*_*A*1_ and all or most of the variability in 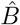 is due to *r*_*B*1_ (e.g., through strong weights for the target channels and small weights for all other channels). Under such circumstances, *Â* and 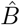 will also be dependent and we have a counterexample to the universal statement *if encoding independence fails, then decoding independence must hold*. One can also choose linear *g_A_* and *g_B_* so that *r*_*A*1_ and *r*_*B*1_ are assigned a weight of zero and have no influence on the decoded variables *Â* and 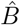. Because in this example all channels in **r**_*A*_ are independent from all channels in **r**_*B*_ except for *r*_*A*1_ and *r*_*B*1_, getting rid of the influence of those two channels over the decoded variables *Â* and 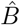 makes them independent (for the same reason that encoding independent implies decoding independence).

Note also that the same assumptions that allowed us to propose that perceptual separability is a form of decoding separability, allow us to conclude that perceptual independence is a form of decoding independence.

### Direct and Indirect Tests of Decoding Separability

#### Direct tests of decoding separability and perceptual separability

Assume that **r** is a vector of neural responses encoding dimension *A* in the brain. If we had access to direct measurements of **r** (e.g., firing rates from single cell recordings or a measure of the activity of a homogeneous neural population), we could use an experimenter-defined decoding function *Â* = *g_E_*(**r**) to estimate dimensional values. Obtaining a large number of decoded values *Â* for each stimulus *A_i_B_j_* allows one to obtain a kernel density estimate (KDE) of *p*(*Â|A_i_B_j_, θ*), represented by 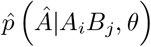. Comparison of such KDEs constitutes a direct test of decoding separability (Equation 14).

Because perceptual separability is a form of decoding separability (Equation 18), the same procedure can be used to obtain the first available direct test of perceptual separability, when a number of conditions are met. First, the vector **r** should include all neural responses encoding dimension *A* in the brain. Second, for all values of **r**, *g_E_*(**r**) = *g_R_*(**r**), so that *g_E_*(**r**) = *x* (each experimentally-decoded value is equal to the perceptual effect). As the vector **r** cannot be identified and measured using currently available methods and *g_R_*(**r**) is unknown, both assumptions appear very difficult to meet.

#### Indirect tests of decoding separability from neuroimaging data

The relations between encoding separability and decoding separability summarized in Figure 3 hold for any decoder, but it can be shown that using a linear decoder allows for a valid test of decoding separability even when indirect measures of neural activity contaminated with measurement error are used, as is the case with fMRI data.

We have assumed the the channel output *r_c_* represents neural activity in a single neuron or a group of neurons with similar properties (e.g., same tuning). Often we do not have access to such direct recordings; rather, we obtain indirect measures of neural activity, which are some function of the activity of several different neural channels. Let *a_m_* represent an indirect measure of neural activity, where *m* = 1, 2, …*M* indexes different instances of the same type of measure (e.g., different voxels in an fMRI experiment or electrodes in an EEG experiment). The measures can be represented by a vector **a** = [*a*_1_,*a*_2_,…*a_M_*], which is a function of the activity of all channels in the encoding model:

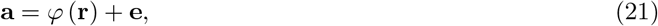

where **e** is a random vector representing measurement error:

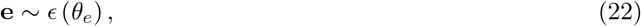

*ϵ* denotes the probability distribution of measurement error, which depends on a set of parameters *θ_e_*. Together, Equations 21 and 22 describe the *measurement model* for **a**.

In a typical multivariate analysis of neuroimaging data, we decode an estimate of a dimensional value *Â* directly from a. We can choose to use a linear decoder for this task, so that

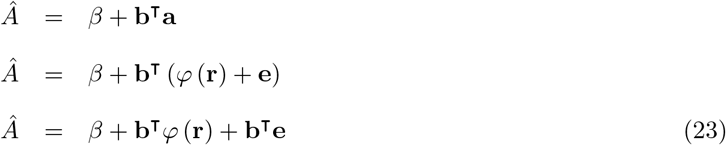

We can think of the estimate *Â* as the sum of two independent random variables: *Â*_**r**_ = *β* + **b^T^***φ*(**r**), which depends exclusively on the distribution of **r**, and *Â_e_* = **b^T^e**, which depends exclusively on the error distribution from Equation 22. The variable *Â*_**r**_ depends on **r** through a composite function. We can think of this composite function as a decoder: *g*(**r**) = *β* + **b***φ*(**r**) and use *p*(*Â*_**r**_|*A_i_B_j_, θ*) to test for decoding separability. Unfortunately, our measurements are contaminated by the variable *Â_e_* with distribution *p*(*Â_e_|θ_e_*). Because *Â* is the sum of two independent random variables, the distribution of *Â* is a convolution of the distribution of each of its components:

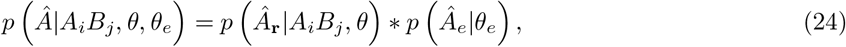

where * denotes the convolution integral.

Thus, KDEs obtained from *Â* decoded from neuroimaging data reflect the target decoding distribution convolved with an error distribution. This means that obtaining direct estimates of GRT perceptual distributions from neuroimaging data may not be possible. Still, it is possible to obtain a valid measure of violations of decoding separability.

Without loss of generality, suppose that we want to measure differences between the distributions *p*(*Â*_**r**_|*A*_1_*B*_1_, *θ*) and *p*(*Â*_**r**_|*A*_1_*B*_2_, *θ*). A number of measures of the distance between two probability densities (such as the *L*1, *L*2 and *L*∞ distances, see [49]) start by computing a difference function:

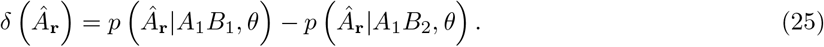

From neuroimaging data, we obtain estimates of the distributions *p*(*Â*_**r**_|*A*_1_*B*_1_, *θ*)\**p*(*Â_e_|θ_e_*) and *p*(*Â*_**r**_|*A*_1_*B*_2_, *θ*)\**p*(*Â_e_|θ_e_*), where we have assumed that the measurement error model does not change with the value of the stimulus in dimension *B*. The difference function between these two distributions is:

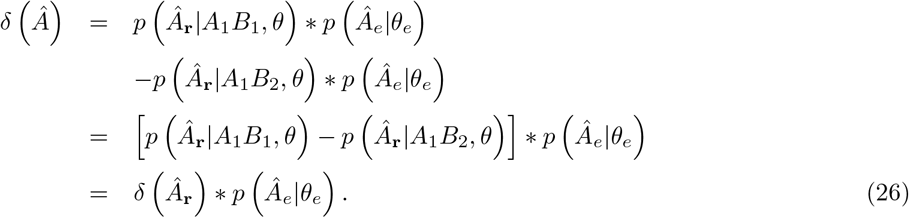

Thus, the difference between noisy KDEs 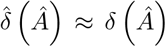 is an estimate of the target difference function *δ*(*Â*_**r**_) convolved with the error kernel *p*(*Â_e_|θ_e_*). Note first that if decoding separability holds, then *δ*(*Â*_**r**_) = 0 and we expect 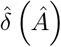 to approximate zero for all values of *Â* as sample size increases. Any deviations from a constant zero function indicate violations of decoding separability. If decoding separability does not hold and *δ*(*Â*_**r**_) ≠ 0 for some *Â*_**r**_, then the shape of the error kernel determines how it affects *δ*(*Â*_**r**_). Under the common assumption that measurement error e is Gaussian with zero mean and covariance matrix Σ_*e*_, *p*(*Â_e_|θ_e_*) will also be Gaussian with zero mean and variance **b^T^**Σ_*e*_**b**. In this case, the convolution attenuates high-frequency fluctuations in the difference function *δ*(*Â*_**r**_). In general, the difference 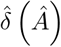 will capture some deviations from decoding separability, but not necessarily all of them.

In sum, as the number of data points used to obtain KDEs increases, a distance measure based on the function 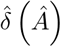 will be approximately zero when there are no violations of decoding separability, and any non-zero value will be the consequence of a violation of decoding separability. This makes such a measure a valid indicator of violations of decoding separability. One measure based on 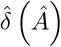 is the L1 norm:

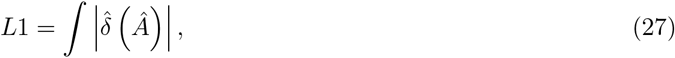

which is the basis for the statistic that we use in our test to measure deviations from decoding separability (DDS statistic; see Materials and Methods section).

Note that the property described by equation 26 holds for distance measures based on the simple difference function 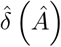. Other commonly-used measures of the distance between two distributions, such as the Kullback-Leibler divergence, are not influenced by measurement error in this straightforward manner, and thus their interpretation is more difficult.

Linear decoders, which are necessary to obtain a valid indirect test of decoding separability, are also the most widely used in the MVPA literature [47]. This allows us to link our framework to this line of research in neuroimaging.

### Relation to previous operational definitions of neural independence

#### Orthogonality of neural representations

Suppose that there are two vectors in the space of encoding channels, *ρ_A_* and *ρ_B_*, representing important summary statistics of how *A* and *B* are encoded, respectively. We can interpret *ρ_A_* as an estimate of the direction along which dimension *A* is encoded, and *ρ_B_* as an estimate of the direction along which dimension *B* is encoded. We can define *encoding vector orthogonality* as the situation in which these two vectors are orthogonal from each other:

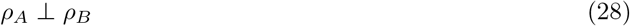

Each choice of statistic *ρ_A_* and *ρ_B_* produces a different form of encoding vector orthogonality. For example, one might be interested in two mean vectors summarizing the encoding distributions for stimuli along the dimensions *A* and *B*. In that case, a simple possibility would be to use stimuli without any value in dimension *A*, represented by *A*_0_, and stimuli without any value in dimension *B*, represented by *B*_0_. Thus, *A_i_B*_0_ would represent a stimulus with a value in dimension *A* only, and *A*_0_*B_j_* would represent a stimulus with a value in dimension *B* only. We might be interested on the mean response of encoding channels to such stimuli, so that *ρ_A_* = **f**(*A_i_B*_0_) and *ρ_B_* = **f**(*A*_0_*B_j_*), or perhaps on the average responses when any level of the dimensions is presented, so that 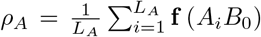 and 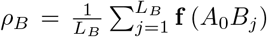. Of course, other possibilities exist (e.g., computing expectations on marginal distributions), and in each case Equation 28 offers a different definition of encoding vector orthogonality.

Linear decoders (Equation 12) also provide estimates of directions in space along which dimensions vary. We might obtain one of such decoders for each dimension, each with its own weight vector **b**, so that *ρ_A_* = **b**_*A*_ and *ρ_B_* = **b**_*B*_. Again, different decoders provide different definitions of encoding vector orthogonality. Note that this is a form of *encoding* vector orthogonality, as the weights are defined in the space of the encoding channels (i.e., 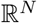) rather than the decoded variables *Â* and 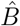. Decoding vector orthogonality cannot be defined, as the decoded variables are scalars.

Neuroscientists have shown great interest in measuring different forms of measurement vector orthogonality, although in many cases they use indirect tests. That is, experimenters usually test the orthogonality of two vectors, **h**_*A*_ and **h**_*B*_ (summarizing information about dimensions *A* and *B*, respectively), defined in a *measurement space* that is a transformation of the original neural *encoding space*. We represent this transformation with *φ_h_*(), to highlight its relation to the measurement model defined in Equation 21. The transformation *φ_h_*() may be known. For example, Kayaert et al. [50] submitted patterns of neural firing rates to multidimensional scaling, and tested the orthogonality of two vectors in the solution space, each representing changes in a dimension of interest. In many cases, however, *φ_h_*() is unknown. This is the case when the test is carried out in a space of indirect activity measures from neuroimaging, as represented by Equation 21. For example, Hadj-Bouziane et al. [6] tested the orthogonality of two vectors of unidimensional fMRI contrasts (one for faces > objects and one for expressive face > neutral face). Another possibility would be to obtain weight vectors representing directions that separate classes best in such measurement space, as seen in Figure 8. Although vectors of unidimensional contrasts (like those used by Hadj Bouziane et al. [6]) and weight vectors do not have the same interpretation [51] (in most cases, they will be completely different vectors), they are both defined within the same measurement space of fMRI voxels or EEG channels. In other cases, the test is carried out in the physical space of the measurements. For example, Baumann et al. [52] estimated **h**_*A*_ and **h**_*B*_ as directions in the physical space of a brain region (the inferior colliculus) along which two dimensions of sound were encoded. In this case, although *φ_h_*() is unknown, the measurement space itself might be interesting and studying it could result in a better understanding of brain function.

More generally, we can define measurement vector orthogonality as the case in which the following relation holds:

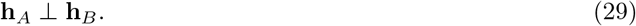

Note that Equation 29 holds if **h**_*A*_ and **h**_*B*_ are mean-centered and their Pearson correlation is zero, as the Pearson correlation of two mean-centered vectors equals the cosine of their angle. Previous researchers have used both the angle between vectors [52, 50] and their correlation [6] as measures of orthogonality.

Many researchers seem to make the implicit assumption that the measurement space (e.g., the estimated activity patterns in an fMRI study) can be directly interpreted as the encoding space. In that case, different choices for **h**_*A*_ and **h**_*B*_ have widely different interpretations (e.g., contrast vectors versus classifier weights; see [51]). However, we believe that this assumption is untenable in general, and particularly difficult to justify in the case of neuroimaging, where the transformation *φ_h_*() is known to involve a series of complicated physical and biological processes.

We suspect that neuroscientists apply the different tests in the hope of learning something about the underlying neural representations at the level of encoding. To understand whether and how that goal can be achieved, the important issue is not so much what version of the test is used or what form of encoding vector orthogonality one is interested in. Rather, the important question is what can be inferred about encoding vector orthogonality in general from the results of any indirect test. That is, given a choice of *ρ_A_* and *ρ_B_* as encoding vectors of interest, a choice of **h**_*A*_ and **h**_*B*_ as measurement vectors of interest, and a particular transformation *φ_h_*() from the encoding space to the measurement space: What does observing measurement vector orthogonality (Equation 29) tell us about encoding vector orthogonality (Equation 28). The answer is: in most cases, nothing, even when one makes the simplifying assumption that **h**_*A*_ = *φ_h_*(*ρ_A_*) and **h**_*B*_ = *φ_h_*(*ρ_B_*).

The reason is that encoding vector orthogonality is defined as a 90-degree angle between *ρ_A_* and *ρ_B_* (Equation 28), and only rigid transformations (i.e., not even all linear transformations) can preserve this angle. Thus, when *φ_h_*() is unknown and involves more than rigid transformations (the most common case), a 90-degree angle between **h**_*A*_ and **h**_*B*_ could be accompanied by any angle value between *ρ_A_* and *ρ_B_*, depending on the specifics of the unknown *φ_h_*(). The test might be uninformative even in the rare case in which *φ_h_*() is known and linear, as computing the angle between *ρ_A_* and *ρ_B_* from **h**_*A*_ and **h**_*B*_ would require for *φ_h_*() to have an inverse. In all the examples from the literature discussed earlier, an observation of measurement vector orthogonality does not provide any information about encoding vector orthogonality. However, as indicated above, the test might provide useful information about other aspects of brain representation different from encoding, as is the case when the measurement space of **h**_*A*_ and **h**_*B*_ is itself interesting (e.g., physical space in studies of functional brain topography, see [52]).

It is possible to link a specific form of encoding vector orthogonality to the concept of perceptual independence (equation 2) from GRT. When stimulus *A_i_B_j_* is presented, it is represented as a new vector **r** in the same encoding space that contains *ρ_A_* and *ρ_B_*. If we assume that (i) the projection of this vector onto *ρ_A_* and *ρ_B_* corresponds to the perceived values of dimensions *A* and *B*, then measurement vector orthogonality is equivalent to *dimensional orthogonality* [53, 54]. One case in which this assumption is met is when the readout functions from Equation 17 (i.e., the functions used by readout neurons to decode perceptual effects from neural activities) are linear. In that case, the weight vectors of the linear decoders correspond to *ρ_A_* and *ρ_B_* and their orthogonality is equivalent to orthogonality of the perceptual dimensions within the encoding space. Ashby and Townsend [14] showed that if, in addition, (ii) the trial-by-trial perceptual effects have a multivariate Gaussian distribution, 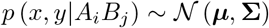, and (iii) Σ does not depend on the stimulus (i.e., all perceptual distributions have identical variance-covariance matrices), then dimensional orthogonality and perceptual independence (as defined in Equation 2) are equivalent.

Assumptions (i)-(iii) also allow one to link measurement vector orthogonality and perceptual independence. However, in this case the assumptions seem extremely strong and hard to meet. Although assumptions (ii) and (iii) are common in the psychophysics literature and they may be justifiable, assumption (i) is very problematic, as there seems to be no way to guarantee that the estimates hA and hB must correspond to perceived stimulus dimensions. In addition, note that dimensional orthogonality must be defined in a particular space of interest. Determining whether the two perceptual dimensions are orthogonal within the original encoding space seems like an interesting question worth pursuing. On the other hand, determining whether the two perceptual dimensions are orthogonal within some arbitrary measurement space does not carry the same weight.

Finally, the problems with the measurement vector orthogonality test are not restricted to their difficult interpretation. In addition, there are practical issues with the way in which the tests are applied. In particular, orthogonality tests are best suited to provide evidence of violations of orthogonality, but they are usually applied to provide evidence of its presence. More specifically, if **h**_*A*_ and **h**_*B*_ were random vectors, then one would expect their correlation to be close to zero (i.e., orthogonal vectors), especially for highdimensional vectors such as those studied by Hadj-Bouziane et al. [6]. Under such circumstances, a finding of orthogonality is expected even from completely random data. In addition, orthogonality corresponds to a single value (zero correlation or 90-degree angle) and therefore evidence of orthogonality requires special statistical tests that can provide evidence for that specific value (e.g., evidence for the null in a Bayes factor test, or a small confidence interval containing the target value). Such tests have not been used in previous tests of orthogonality. Here, we will explore whether it is possible to find *violations* of orthogonality in our data, rather than trying to find evidence *for* orthogonality as in previous studies.

In sum, measurement vector orthogonality is an operational test of independence of neural representations, and several researchers have used some version of it in past studies. In general, the results of this test cannot be related to a corresponding property of stimulus encoding, which we named encoding vector orthogonality. If some strong assumptions are met, the test can be related to the concept of perceptual independence from GRT, which is conceptually distinct from the several forms of separability on which we have focused here. In particular, perceptual independence is a property of a single stimulus. It holds if different stimulus components are processed independently of each other. In contrast, separability is a property of an ensemble of stimuli. It holds if processing of one component is unaffected by changes in other components. In practical terms, we would expect that violations of measurement representation orthogonality would be unrelated to violations of decoding and encoding separability, as they measure completely different concepts.

#### Classification accuracy invariance and generalization

A second operational test of independence of neural representations, more closely related to the separability measures investigated in this article, has been recently used in research on invariance of face representation. In this test, activity patterns in a given brain region are classified according to some target dimension. For example, activity patterns in visual cortex could be classified according to the identity of faces presented during an experiment. Then the classifier is tested with new patterns, produced during presentations of the same identities but with changes in some irrelevant face dimension, such as viewpoint or expression [55, 56, 57]. If the classifier’s accuracy is significantly above chance, it is concluded that the representations of the target dimension (face identity) are invariant to changes in the irrelevant dimension. A simpler version of the test simply checks for significant classification accuracy using all data [9], but this is much less informative than a test based on generalization after changes in the irrelevant dimension [55].

Formally, let *ℓ_i_* represent a label returned by the classifier indicating that it has estimated that level *i* of dimension *A* has been presented, and suppose the experiment includes a total of *L_A_* different levels of dimension A. Then *classification accuracy generalization* is defined in the following way:

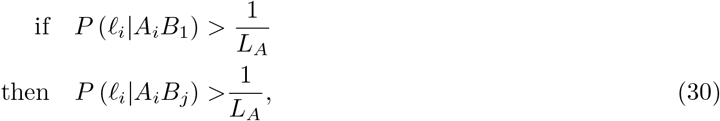

for all *i* and *j*. That is, if the probability of correct classification of the level of *A* is higher than chance 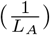 at level 1 of dimension *B*, then it must be higher than chance at all levels of *B* for classification accuracy generalization to hold.

It is possible to indirectly relate classification accuracy generalization to encoding separability, through its clear relation to decoding separability. We do this first for the case in which there is access to direct measures of neural activity that are used to estimate *Â*, as in Equation 11. Because *Â* is a noisy estimate that can assume any value in 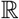, a classifier partitions this space into *L_A_* regions, one for each of the values of dimension *A* included in the experiment. Let 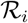 represent the region associated with label *ℓ_i_*, so that the classifier assigns this label to a neural pattern when 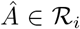. Each 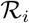 may be a single continuous interval in 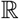 or composed of several such intervals, and the union of all 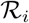 completely covers the real line 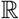. When the decoding distribution is known, classification accuracy for level *i* of dimension *A* is:

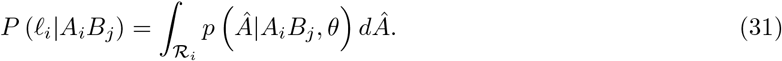

Equation 31 relates classification accuracy to the distribution of decoded values on the target dimension *A*. From this we know that *if decoding separability holds, then classification accuracy generalization must hold*. This is true because when decoding separability holds, the distribution *p*(*Â|A_i_B_j_,θ*) inside the integral in Equation 31 is the same for all values of *j*, *P*(*ℓ_i_|A_i_B_j_*) is therefore the same for all values of *j* and the relation in Equation 30 holds. On the other hand, *if decoding separability fails, then classification generalization may hold or fail*. Without loss of generality, assume that decoding separability fails because *p*(*Â*|*A_i_B*_1_,*θ*) ≠ *p*(*Â|A_i_B*_2_,*θ*). Regardless of the shape of *p*(*Â|A_i_B*_1_,*θ*) and the region covered by 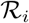, there are an infinite number of other shapes for *p*(*Â|A_i_B*_2_,*θ*) that will preserve the area under the curve inside region 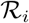 constant, making the value of *P*(*ℓ_i_|A_i_B_j_*) constant across changes in *j*, which ensures that the relation in Equation 30 holds. Alternatively, regardless of the shape of *p*(*Â|A_i_B*_1_,*θ*) and the region covered by 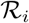, there are also an infinite number of other shapes for *p*(*Â|A_i_B*_2_,*θ*) that will change the area under the curve inside region 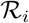, making the value of *P*(*ℓ_i_|A_i_B_j_*) change across changes in *j*. Under such circumstances, the relation in Equation 30 may or may not hold, depending on whether or not the shape of *p*(*Â|A_i_B*_2_,*θ*) produces an area under the curve inside 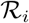 that is larger than 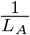.

These considerations point toward an intermediate kind of invariance between decoding separability and classification accuracy generalization, which we call *classification accuracy invariance*, defined as the case in which classification accuracy for levels of dimension *A* is invariant across changes in the stimulus on a second, irrelevant dimension. That is, classification accuracy invariance of dimension *A* with respect to dimension *B* holds if and only if, for all values of *i* and *j*:

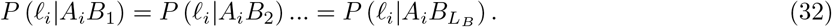

Decoding separability (Equation 14), classification accuracy invariance (Equation 32), and classification accuracy generalization (Equation 30) are related to one another as described in Figure 4. The proofs offered earlier relating decoding separability and classification accuracy generalization already include classification accuracy invariance as an intermediate form of invariance for which the relations in Figure 4 hold.

**Figure 4:**
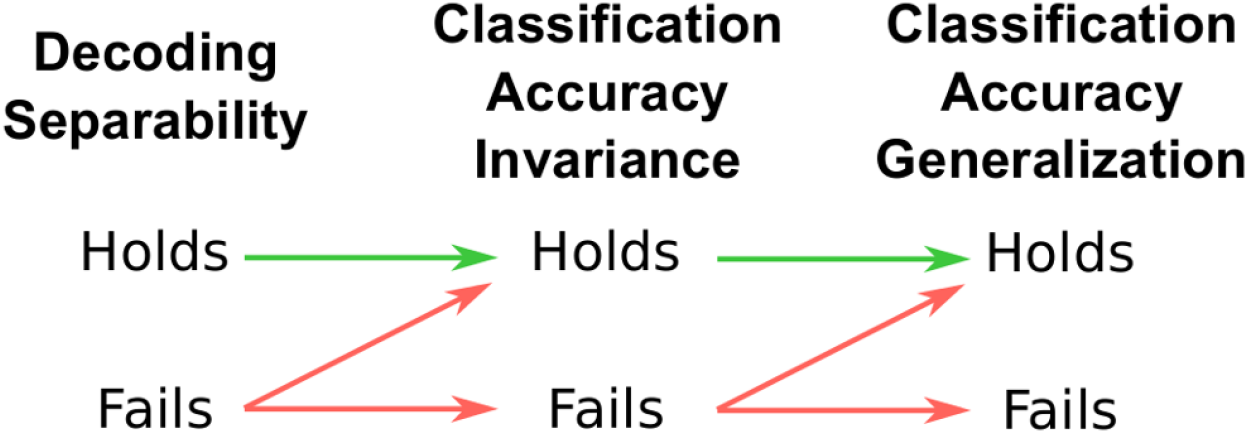
Summary of the relation between decoding separability, classification accuracy invariance and classification accuracy generalization, according to our extension to GRT. Arrows should be interpreted as conditional statements of the form “if X, then Y”.

What happens when classification accuracy invariance and generalization are evaluated through indirect measures of neural activity, such as those obtained from neuroimaging? This is the way in which such tests have been most commonly applied [55, 56, 57, 9]. Remember that in this case, the addition of measurement and noise models (Equations 21 and 22) considerably changes the distribution of *Â* estimates obtained from a linear decoder, which is the result of convolving a distribution of decoded values and the distribution of measurement error. This is likely to change the specific classification accuracies *P*(*ℓ_i_|A_i_B_j_*) but it should not change their relations as defined in Equations 30 and 32, under the assumption that the measurement error model does not change with the value of the stimulus on dimension *B*.

The theoretical results summarized in Figure 4 reveal two issues with the classification accuracy generalization test as it is currently applied in the neuroimaging literature. The first and most important issue is that finding that classification accuracy generalization holds does not provide any information about encoding separability. On the contrary, what provides information about violations of encoding separability is finding a *violation* of classification accuracy generalization. Thus, while this test seems to be valid and useful, it is currently applied and interpreted in the wrong way [55, 56, 57, 9]. It is possible that finding classification accuracy generalization may provide information about other properties of encoding, but such properties are yet to be identified within a formal framework like the one presented here. Another possibility is that classification accuracy generalization could provide information about encoding separability in special circumstances (i.e., for a specific choice of encoding, decoding, measurement and error models), but again such possibilities are yet to be shown. The second issue with the classification accuracy generalization test is that, given the relations shown in Figures 3 and 4, it provides less information about encoding separability than the decoding separability test proposed in the previous section. In Figure 4, each logical step away from decoding separability implies that a number of violations of encoding separability might go undetected, due to the up-diagonal arrow at each step. Thus, the classification accuracy generalization test is likely to be less sensitive to violations of encoding separability than a decoding separability test. If the goal of a study is to learn about encoding separability, then the wiser decision is to focus on a test of decoding separability, rather than on tests of classification accuracy. An aspect of the lack of sensitivity of the classification accuracy generalization test is the fact that it requires accuracies significantly above chance to be applied and thus should always be applied using an optimal classifier. On the other hand, the decoding separability test offers a sensitive measure of deviations from encoding separability regardless of what decoder is used, including situations in which the decoder is not optimal and/or does not achieve significant classification accuracy.

#### Pattern difference invariance

The work of Allefeld and Haynes [58] suggests another way to indirectly test encoding separability in neuroimaging. These authors have proposed the multivariate general linear model (MGLM) as an alternative to decoding for the analysis of the multivariate activity patterns typically measured with neuroimaging. One feature of the MGLM is that it allows researchers to test both main effects and interactions in an experimental design. For example, if we are studying the effect of variations in the level of dimensions *A* and *B* on observed multivariate patterns, *A* main effect might answer the question of how multivariate patterns change with variations in the level of *A*. On the other hand, an interaction effect answers the question of how the pattern difference between different levels of *A* changes with variations in the level of *B*.

Thus, we can define *pattern difference invariance* of dimension *A* with respect to dimension *B* as the case in which the pattern difference produced by changes in the level of *A* does not change with the level of dimension *B*. Within the MGLM framework, failures of pattern difference invariance can be tested through the interaction between factors *A* and *B*. This test has not been applied or proposed for the study of independence of neural representations in the past, but it seems like a straightforward application of the more general MGLM framework advanced by Allefeld and Haynes [58].

It is important to note here that the MGLM approach has been proposed as a way to analyze neuroimaging data, and therefore the pattern difference invariance test would be applied to data that is only indirectly related to the underlying neural activity patterns **r**, and that is contaminated with measurement error. Thus, we must determine whether this is a valid test of properties of encoding such as encoding separability. To simplify our discussion, assume that the parameter vectors estimated by the MGLM represent estimates of the activity patterns a defined within our framework (see measurement model in Equation 21). Under the assumption that the measurement model is the same across changes in the level of the irrelevant dimension, we know that *if encoding separability holds, then pattern difference invariance must hold*, as identical encoding distributions should produce identical distributions of measured activity patterns a. On the other hand, *if encoding separability fails, then pattern difference invariance may hold or fail*. This is easy to prove by counterexamples, as we did previously when linking encoding separability and decoding separability. As before, we assume that dimension *A* is encoded through the model with Gaussian channel noise (Equation 7). We also assume a linear measurement model (Equation 21), which is very common in the neuroimaging literature:

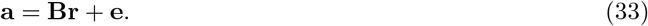

Finally, as before, we suppose that encoding separability fails due to a difference in the tuning functions of *A*_1_*B*_1_ and *A*_1_*B*_2_, whereas the tuning functions of *A*_2_*B*1 and *A*_2_*B*_2_ are identical:

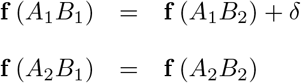

Under the assumptions listed above, the tuning functions only affect the mean of the measured activity patterns. The variance of each measure of neural activity is a linear combination of channel noise variances plus the measurement error variance. Those variances can be ignored, as they are not affected by differences in the tuning functions and, in addition, they are assumed to be identical across stimuli in the MGLM framework. When pattern difference invariance holds, we have that:

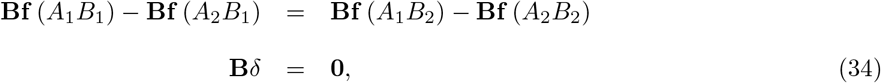

whereas when pattern difference invariance fails, we can show that:

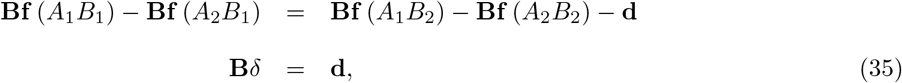

where **d** represents a difference vector. For any given *δ*, equations 34 and 35 both can be satisfied by an infinite number of matrices **B**, yielding a model in which encoding separability fails and pattern difference invariance either holds (Equation 34) or fails (Equation 35).

Thus, the pattern difference invariance test has the same logical relation to encoding separability as decoding separability (Figure 3). Unfortunately, pattern difference invariance is not directly related to decoding separability, as was the case for classification accuracy generalization. Without considerable more work, it is difficult to determine which of the two tests will prove to be more useful in different situations. However, we believe that the decoding separability test proposed earlier is a better choice in most cases, for two reasons. The first reason is that the decoding separability test does not make any assumptions about the multivariate distribution of the data, besides assuming that measurement error is additive. Using a linear decoder ensures that the test is valid (we have used Equation 23 as the starting point to showing that this is the case), but such a decoder can be obtained using methods that do not make strong assumptions about the data distribution, such as support vector machines. On the other hand, a test based on the MGLM makes the strong assumption that the data is multivariate normal with equal variance-covariance matrix across conditions (i.e., stimuli). The second reason is that a test based on the MGLM is based on a comparison between parameter differences, and thus is insensitive to differences between distributions in higher-order moments. When enough data are collected, the decoding separability test is also sensitive to differences in higher-order moments of the decoding distributions (i.e., variance, kurtosis, etc.). Still, the two tests are based on different approaches (one compares activity pattern distributions, the other compares decoding distributions), so only future work will determine their relative usefulness.

An additional theoretical relation between the MGLM framework and GRT can be established. Allefeld and Haynes [58] define *pattern distinctness* as a measure that quantifies the difference between two multivariate activity distributions. When there are only two distributions being compared, the pattern distinctness is related to the Mahalanobis distance between the two activity distributions (e.g., *A*_1_*B*_1_ and *A*_2_*B*_1_). Thomas [59, 60] has shown that in some cases the Mahalanobis distance between two distributions can be related to a generalized *d*′ measure that quantifies sensitivity for multivariate distributions, which provides a theoretical link between the way in which discriminability is measured in the MGLM and multidimensional signal detection theory.

## Summary of Theoretical Results

Here we summarize the previous theoretical results, with an emphasis on how they can be applied to the empirical study of perceptual independence by psychologists and neuroscientists. A summary of all the theoretical relations described in previous sections can be seen in Figure 5.

**Figure 5:**
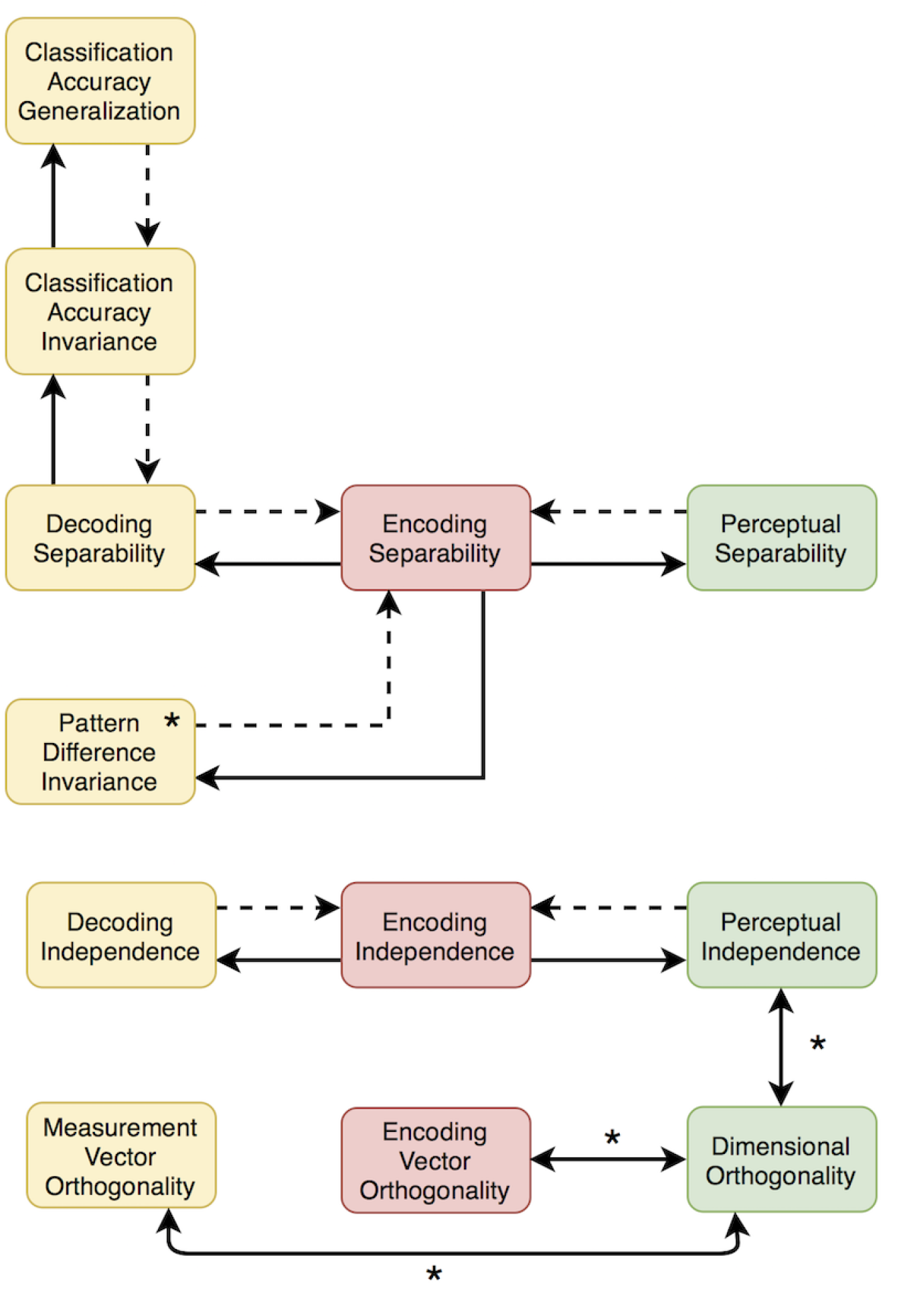
Summary of the theoretical relations found here. Yellow rectangles represent tests that can be applied to neuroimaging data. Red rectangles represent properties of neural encoding. Green rectangles represent properties of perceptual representations. Solid directional arrows indicate that if the concept where the arrow starts is true, then the concept where the arrow ends must be true as well. Dotted directional arrows indicate that if the concept where the arrow starts is false, then the concept where the arrow ends must be false as well. Bidirectional arrows indicate that the two concepts are equivalent. Asterisks are displayed on relations or tests that depend on relatively strong assumptions (see main text for details).

First, because perceptual separability can be considered a form of decoding separability, and due to the relations summarized in Figure 3, any failure of perceptual separability should be reflected in a failure of encoding separability somewhere in the brain. This means that any psychophysical study reporting a failure of perceptual separability provides a hypothesis to be tested by a neuroscientific study: that a corresponding failure of encoding separability should be found, probably in sensory areas thought to encode the target dimension. Second, such neuroscientific studies can be performed using direct measures of neural activity, such as those provided by single-cell recordings or local field potentials, or indirect measures of neural activity contaminated by measurement error, such as those provided by EEG and fMRI. Using traditional linear decoding strategies on indirect measures of neural activity, the decoded dimensional values still offer a basis for a valid test of decoding separability, and any violation of decoding separability found within a given brain region reflects a violation of encoding separability by the neural population in that region. It must be stressed that a failure of encoding separability is a valid inference that can be made from decoding of neuroimaging data, but such data do not provide a basis to make any strong inferences about the presence of encoding separability. A weak inference can be made, based on the lack of evidence for a violation, but this is analogous to accepting the null in a traditional statistical test. A relatively stronger inference of encoding separability could be made on the basis of assumptions about the neuroimaging measurement model, but researchers should clearly identify such assumptions. Our recommendation to researchers is to be cautious about concluding that separability (or “invariance”) holds at the neural level from neuroimaging data, or even from decoding of direct measures of neural activity (e.g. [61]; a related point was made in [62, 63]).

Finally, we have shown that operational tests of independence available in the literature can be formally defined and re-interpreted within the framework presented here. We showed that, when some strong assumptions are met, the measurement vector orthogonality test [52, 6, 50] is related to the concept of perceptual independence from the traditional GRT, but it is unlikely to be related to a corresponding property of stimulus encoding. On the other hand, the classification accuracy generalization test promoted by Anzellotti and Caramazza [57, 55, 56] can lead to valid inferences about encoding separability. However, the way in which the test has been applied might lead to conclusions of invariance or separability that are in general unjustified, unless one is interested in decoding separability only, and not in the separability of underlying brain representations. In addition, the classification accuracy generalization test is likely to provide less information than our decoding separability test. The MGLM approach proposed by Allefeld and Haynes [58] suggests an alternative way to indirectly test encoding separability in neuroimaging. The resulting pattern difference invariance test seems like a valid test of violations of encoding separability, but it is based on strong assumptions about the distribution of the neuroimaging data that are not necessary when the decoding separability test is applied.

### An Application to the Study of Encoding Separability of Face Identity and Expression

Here we present an application of our framework to the study of encoding separability of face identity and expression. This application serves as a way to illustrate the kind of question that this framework can help answer and the concrete steps that researchers should take to apply the framework in their research.

Information about a number of properties can be extracted from a single face, including identity and emotional expression. The influential model of Bruce and Young [3] proposed that these two face dimensions are processed independently, motivating a large number of psychophysical studies aimed at testing this hypothesis [64, 33, 65, 66, 34, 67, 68, 69, 35, 13, 70, 71, 72]. Neurobiological theories of visual face processing [4, 73] also propose relatively independent processing of face emotion and identity, through anatomically and functionally differentiated pathways. A ventral pathway projecting from the occipital face area (OFA) to the fusiform face area (FFA) would mediate the processing of invariant aspects of faces, such as identity. A dorsal pathway projecting from the OFA to the posterior superior temporal sulcus (pSTS) would mediate the processing of changeable aspects of faces, such as emotional expression. Recent reviews [74, 75] conclude that the two pathways are indeed relatively separated and functionally differentiated, with the ventral pathway being involved in the representation of face form information–including invariant aspects of face shape such as identity–, and the dorsal pathway involved in the representation of face motion information–including rapidly changeable aspects of faces such as expression. According to this revised framework, both identity and expression information may be encoded in either pathway, but exactly what information about each dimension is encoded would differ between pathways.

The psychophysical and neurobiological lines of research in this area have remained relatively independent across the years, with no attempt to integrate results across levels of analysis despite the similarity of the central questions guiding their research. In addition, both lines have relied largely on operational definitions of independence that, while having face validity, are usually not linked to any theoretical definition. As indicated in the introduction, this approach makes it difficult to interpret contradictory results.

Thus, the study of independence of face identity and expression is a particularly good example of an area in which our extended GRT framework can provide helpful research tools. Our theory can provide a much-needed theoretical integration across levels of analysis and tests, as well as more rigorous definitions of independence and ways to measure it. We have recently performed a GRT analysis of psychophysical data to study the perceptual separability of identity and expression [13]. The results suggested that, for the stimuli used in that study and after accounting for decisional factors, emotional expression was perceptually separable from identity, but identity was not perceptually separable from emotional expression. From these results, our current framework (see Figure 3) predicts that encoding separability of identity from expression must fail somewhere in the areas representing face information, and that we should be able to find evidence of failures of decoding separability in those areas. The predictions regarding encoding separability of emotional expression are less straightforward: as there are no violations of perceptual separability in the behavioral data, violations of encoding separability seem unlikely, but are still possible.

Here, we acquired fMRI data from participants while they looked at the same stimuli and completed the same task as in our previous psychophysical study (see Materials and Methods). The stimuli were images of four faces, which resulted from two different male identities showing two different emotional expressions (neutral and sad). Participants performed a simple stimulus identification task. In each trial, a single stimulus was flashed in the screen and the participant had to identify the specific combination of identity and emotional expression that had been shown. This required participants to pay attention to both identity and expression to attain good performance. The participants received feedback about the correctness of their responses. The task was given to participants in runs that lasted around 10 minutes, during which each image was repeated 25 times. Participants completed three of such runs, and in addition they completed a standard functional localizer run [76] that allowed to obtain the approximate location of face-related regions.

Performance in the task during the scanning session was high, with a mean of 81.67% (SE = 5.18%). Single-trial estimates of stimulus-related activity were used as input to the decoding separability test described earlier. Because we did not have specific hypotheses about the location of areas showing failures of encoding separability, we performed a whole-brain searchlight analysis [77], to determine which small circular regions (radius of 3 voxels) showed violations of decoding separability, and therefore violations of encoding separability. To spatially localize violations of encoding separability relative to areas in the face network, we found such areas with the help of a standard functional localizer.

The results from this analysis did not reveal any significant violations of decoding separability, either for identity or emotion. Further exploration revealed that our standardized DDS index was consistently negative in the full-brain maps, suggesting that our method of standardization might have produced an index that was too conservative. That is, the *DDS* was standardized to represent a percentile value (ranging from 0 to 100) re-centered around the middle of the distribution (i.e., ranging from -50 to 50). Under the null hypothesis of decoding separability, the distribution of this DDS would be driven only by noise in the data, and we would expect the standardized measure to hover around zero, with similar areas of the brain maps being positive and negative. On the other hand, we would not expect values consistently lower than zero, as this would mean that the estimated decoding distributions are consistently more similar to one another than expected under decoding separability. As the expectation under decoding separability is that the distributions are identical, this seemed like a problem. We reasoned that one solution would be to use the difference in DDS index between the identity and emotion analyses as the main test statistic, to allow one map to serve as control for the other. We must underscore that this is an exploratory analysis, and its results should be confirmed by future research. Increasing power with a larger sample (either a larger number of participants or a larger number of trials) would be helpful to obtain reliable results with a conservative test. Figure 6 shows the main results of this analysis, displayed over a flat cortical map. Face-selective areas found through the functional localizer are outlined in the figure. Outlined in green are face-selective areas showing higher activity during the presentation of faces than during the presentation of other objects. Outlined in red are areas showing higher activity during the presentation of emotional faces than during the presentation of neutral faces. The figure also shows clusters of significant violations of decoding separability, depicted in red-yellow for the identity > emotion contrast. A single large cluster (483 2mm voxels) was found to be significant, covering parts of the left STS and superior temporal gyrus (peak location in MNI coordinates: -60, -14, 2). This cluster only slightly overlapped with an area of the face network in the pSTS (green contour). No significant violations were found for the emotion > identity contrast.

**Figure 6:**
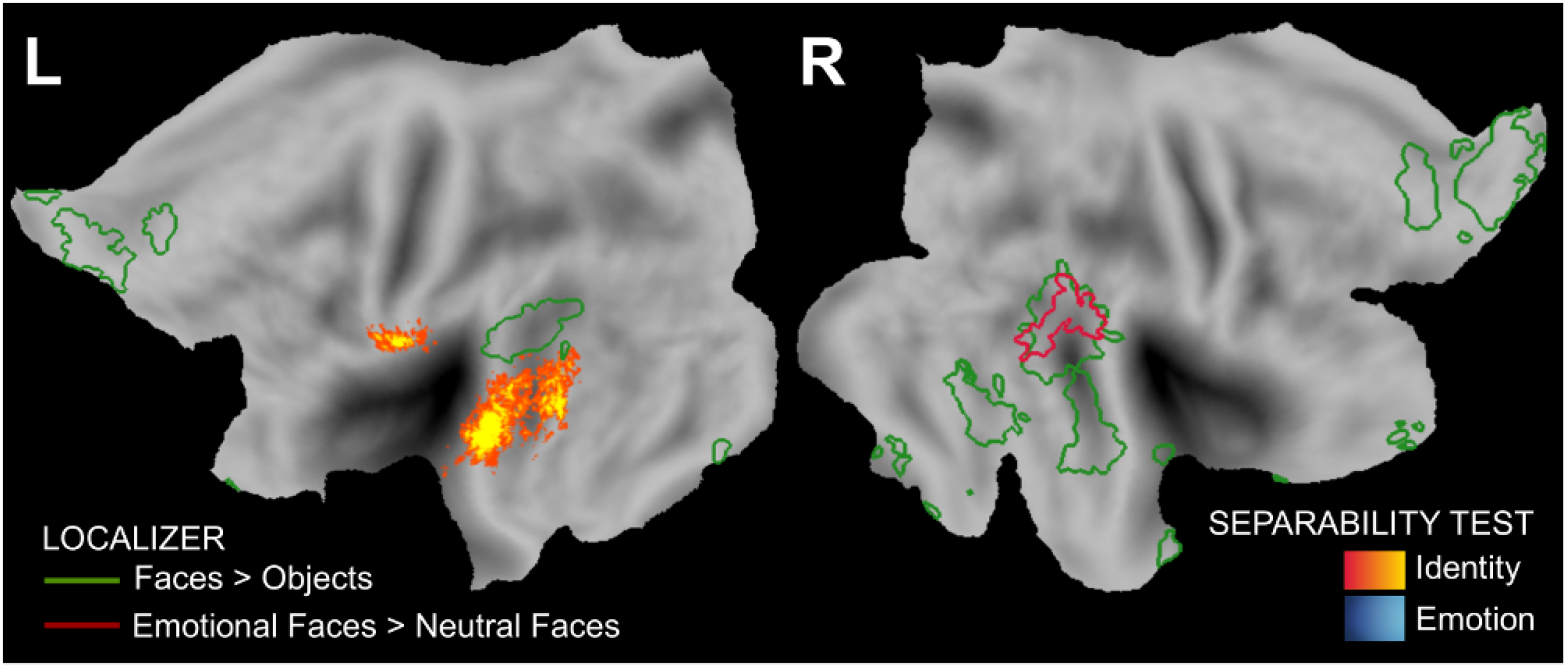
Results of the searchlight decoding separability test. Yellow-red clusters represent regions in which violations of decoding separability of identity were stronger than violations of decoding separability of emotional expression. There were no regions in which violations of decoding separability of emotional expression were stronger than violations of decoding separability of identity. Green and red lines delimit face areas from the functional localizer.

The results shown in Figure 6 are in line with the previous psychophysical results [13] and the relations depicted in Figure 3, as they provide evidence of stronger violations of decoding separability for identity than for emotional expression, but not the other way around. This asymmetry in the separability of neural representations is analogous to the asymmetry in perceptual separability found in our previous psychophysical study, and thus makes intuitive sense. Although this asymmetry was not a strong prediction from the theory (which simply predicts violations of decoding separability for identity, but is ambiguous about violations of decoding separability for emotional expression), it suggests that there is at least an empirical correspondence between asymmetries of separability in perceptual and brain representations.

### Comparison with measurement vector orthogonality

We implemented a version of the measurement vector orthogonality test [52, 50] discussed earlier. At each searchlight, we measured representation orthogonality by correlating the weights from the classifier used in the previous analyses of identity and emotion. A correlation of zero is equivalent to orthogonality of the two weight vectors, and therefore any deviation from a zero correlation is indicative of a violation of measurement vector orthogonality. As mentioned earlier, finding such violations of orthogonality is more informative than finding evidence for orthogonality. The resulting individual orthogonality maps were submitted to the same permutation test previously used for separability maps. No violations of measurement vector orthogonality were found in this analysis.

Note that this finding of measurement vector orthogonality has been taken by other researchers to mean that information about face identity and emotional expression is represented independently in the visual system [6]. Our theoretical results allow us to draw a different conclusion: measurement vector orthogonality cannot be easily linked to a corresponding property of stimulus encoding, as even a linear transformation from the space of the encoding model to the space of indirect measures of neural activity does not necessarily preserve angles.

From the point of view of GRT, decoding separability and measurement vector orthogonality seem to measure unrelated properties of perception and neural encoding. From the point of view of perception, decoding separability is related to the GRT concept of perceptual separability, whereas measurement vector orthogonality is related to the GRT concept of perceptual independence. In both cases, however, the tests are related to the corresponding GRT concepts through a number of strong assumptions. From the point of view of encoding, decoding separability is related to the concept of encoding separability, whereas measurement vector orthogonality seems difficult to relate to any property of encoding. For these reasons, we expected the magnitude of violations of orthogonality and violations of separability to be unrelated. To test this hypothesis, we took the group statistical maps obtained from the permutation test in the analysis of decoding separability and computed their Pearson correlation with the corresponding maps from the current analysis of representation orthogonality. There was a small but significant correlation between the orthogonality map and both the map of deviations of separability for identity, *r*=−0.1162 (*p*<0.0001), and the map of deviations of separability for emotion, *r*=0.0666 (*p*<0.0001). These correlations are significant due to the large number of voxels used to calculate them, but their magnitudes are very small. With these correlations, only 1.35% of the variability in the separability map for identity and 0.44% of the variability in the separability map for expression can be explained by variability in the orthogonality map. Still, the fact that the correlations are significant in real data is important, and some unknown relation between measurement vector orthogonality and decoding separability may underlie these results. Future theoretical work will be necessary to clarify these points.

### ROI-based decoding separability test

An additional ROI-based analysis was performed, with three goals in mind. First, we wanted to determine whether directly testing face-selective regions would result in some evidence of violations of decoding separability, as the only cluster showing such deviations in the searchlight analysis overlapped very little with face-selective regions from the localizer (see Figure 6). Second, we wanted to more clearly determine whether there are meaningful variations in the amount of separability between different regions. Finally, we wanted to explore the behavior of our decoding separability analysis in control regions. The included ROIs are face-selective areas (OFA, FFA, STS) and two control regions: V1, which is known to be sensitive to low-level visual features and thus might show deviations of decoding separability (any change in the faces would produce changes in low-level features), and the lateral ventricles, which give us information about the behavior of our statistic when there is very little underlying signal. Some information may be available at the ventricle ROIs that is leaked from adjacent regions, but we would expect that here our statistic should show decoding separability, as the decoding distributions should be almost completely determined by measurement noise.

Results are shown in Figure 7. Figure 7 shows mean standardized DDS values across all ROIs included in the analysis, with error bars representing standard errors of the mean. The scale of the statistic displayed in Figure 7 is different from that of the full-brain maps, as proportions rather than percentiles are used and the median has not been subtracted. We tested whether any of these means was significantly higher than the value of 0.5 expected under decoding separability through *t*-tests (directional, uncorrected). The only ROIs showing significant violations of decoding separability were the left V1 in the analysis of identity, *t*(15)=2.04, *p*<.05, and the right STS in the analysis of emotion, *t*(20)=2.05, *p*<.05. Due to the large number of tests and the fact that our experiment was not originally designed with ROI-based analyses in mind, none of the tests is significant after the application of a correction for multiple comparisons. For this reason, the results shown in Figure 7 should be taken as only suggestive and exploratory. Still, the evidence is encouraging as it suggests that: (1) deviations from decoding separability are not significant in the control areas assumed to include mostly measurement noise (left and right ventricles), (2) deviations from decoding separability are significant in one of the control areas thought to involve such deviations (left V1), and (3) deviations from separability were very low across face-selective areas, with the exception of the right STS, which showed a significant deviation from decoding separability of emotion from identity. Also note that, for most face-selective regions, the mean DDS is consistently below the value of 0.5 which, as mentioned earlier, suggests that the DDS is a conservative measure of failures of decoding separability, at least in areas thought to encode the dimensions under study.

**Figure 7:**
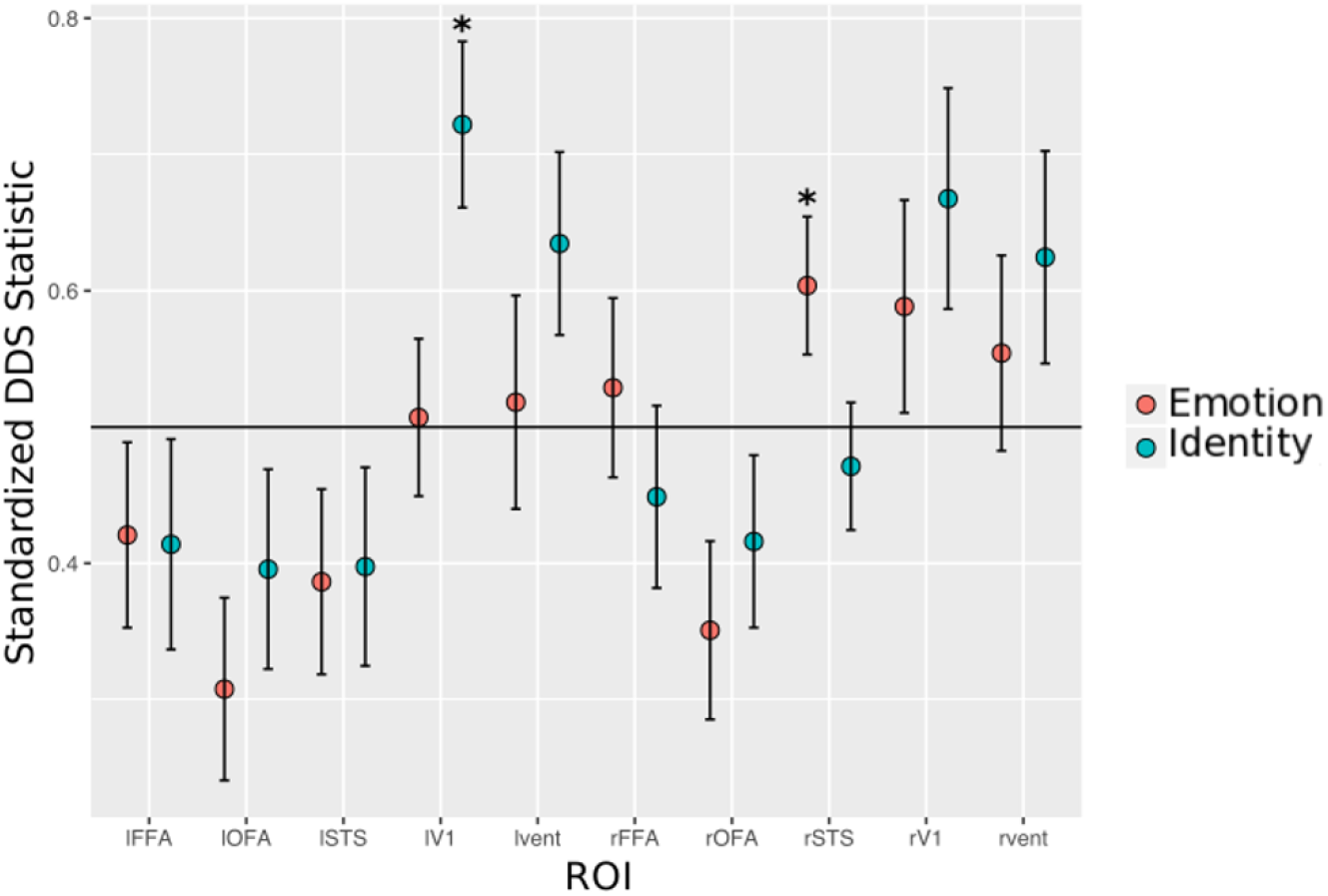
Results of the ROI-based decoding separability test. The y-axis reports the standardized deviations from decoding separability (DDS) statistic. The points represent mean values and the error bars represent standard error of the mean. When decoding separability holds, this index should have a value around 0.5, which is represented with a horizontal dotted line. Mean statistics that were found to be significant (t-test, uncorrected) are marked with an asterisk.

## Discussion

Here we have linked multidimensional signal detection theory from psychophysics and encoding models from computational neuroscience within a single theoretical framework. This allowed us, for the first time, to link the results from psychophysical and neurobiological studies aimed at determining independent processing of stimulus properties. Unlike previous approaches, our framework formally specifies the relation between behavioral and neural tests of separability, providing the tools for a truly integrative research approach in the study of independence.

In the past, neuroimaging studies have been limited to a choice between decoding and encoding approaches to data analysis [29]. Decoding approaches focus on answering questions about *what* is encoded in a given brain region, while making no assumptions about *how* exactly that information is encoded; this lack of commitment to an encoding model is both their strength, as they provide useful results regardless of how information is encoded, and their weakness, as they are limited regarding what kind of question they can answer. In contrast, encoding approaches focus on answering questions about *how* a specific stimulus property is encoded in a brain region, but they do this by assuming the encoding model and determining whether it can help to accurately predict data. Their weakness is that there are a very large number of models that could be tested, and no way of knowing a priori whether the best model is included in the analysis. The theory presented here allowed us to identify some properties of encoding models (i.e., encoding separability) that can be inferred from the results of a decoding study. We hope that future theoretical research in this line will allow researchers to link other properties of encoding models to the results of decoding tests, and more generally to the results of any analysis involving measures that are some transformation of the underlying neural activity, as is the case in fMRI and psychophysics.

Although we focused on developing a decoding separability test, the GRT framework presented here is useful to understand the results of other tests of independence as well [5, 4, 11, 6, 7, 9, 57, 10, 12, 8]. Here, we re-interpreted two operational tests of independence previously applied in the literature within our extended GRT framework. We showed that, when some strong assumptions are met, the measurement vector orthogonality test [52, 6, 50] is related to the concept of perceptual independence from the traditional GRT, but it is unlikely to be related to a corresponding property of stimulus encoding. On the other hand, a test based on generalization of classification accuracy [57, 55, 56] can provide information about encoding separability. However, the test is likely to provide less information than a decoding separability test and it has been applied incorrectly, yielding conclusions of separability (invariance) that are in general unjustified. Application of our framework to additional operational tests may require the development of models linking neural activity to the specific measurements made in each test.

The framework and test proposed here are applicable not only to fMRI data, but also to the analysis of single-cell recordings, LFPs, EEG and MEG. This breadth of scope across operational definitions and levels of analysis (single neurons, neural populations at many scales, perception, and behavior), which is rarely seen in neuroscience, is a very important contribution of the present work.

We applied our new framework to the study of independent representation of face identity and emotional expression. Previous research found that, for the set of stimuli studied here, identity is not perceptually separable from emotional expression, whereas emotional expression is perceptually separable from identity [13]. Our results revealed that such lack of perceptual separability is reflected in stronger violations of decoding separability for identity than for emotion in the left temporal cortex, but no stronger violations of decoding separability for emotion than for identity in any brain region.

Several previous fMRI studies have explored the question of whether emotional expression and identity are represented independently in the brain, and an important question is what value is added by a study based on our extended GRT. We believe that our framework provides at least two advantages. The first advantage is the provision of clear links between the results of neuroimaging and behavioral studies using the same stimuli. No previous study could directly link behavior to neural representation in a meaningful way, as in our application of the extended GRT framework. We started our study with clear hypotheses about how fMRI results should reflect the behavioral results, which is preferable to the approach of linking neural and behavioral results through post-hoc theorizing. The second advantage offered by our framework is that it improves our ability to interpret new results in the light of previous results. For example, our data suggest violations of decoding separability of identity from emotion. This does not contradict previous reports of orthogonality of neural representations [6], as we know that decoding separability and measurement vector orthogonality measure different concepts. Researchers have also found that emotion can be decoded from areas linked to processing of identity [9, 78], and identity can be decoded from areas linked to processing of emotion [79]. The issue of whether or not a particular kind of information can be decoded from a brain region is orthogonal to the issue of whether or not it shows encoding separability. Accurate decoding from a particular area indicates that information about a dimension is present in that area but, as indicated earlier, decoding methods are agnostic as to how that information is encoded. The same is true about inaccurate decoding from a particular area. On the other hand, an example of a test that is related to encoding separability is the classification accuracy generalization test of Anzellotti and Caramazza [55, 57, 79]. However, this test has not been applied to the study of independence of identity and emotional expression, but rather to the study of identity across changes in viewpoint and modality.

### Non-linear decoders and measurement models

We have shown that a decoding separability test operating on indirect measures of neural activity can validly detect violations of encoding separability, but one of the conditions in our treatment of this issue was using a linear decoder. When a linear decoder is used, the relation between the target difference between decoding distributions 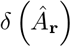 and the measured difference 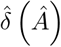 is straightforward (see equation 26), which allows us to know exactly what violations of decoding separability the test can and cannot detect. We believe that the use of a linear decoder is a reasonable requirement for the test, as they are easier to use and interpret than non-linear decoders, and decoding studies in neuroimaging have almost exclusively used linear decoders. However, one open question is to what extent using a different decoder might change the test’s ability to detect specific violations of encoding separability. That is, perhaps a specific type of violation of encoding separability is hidden by a linear decoder, but shown by a non-linear decoder, or vice-versa. This question requires considerable additional theoretical and simulation work. However, we do know that, regardless of what type of decoder is used, if encoding separability holds, then decoding separability must necessarily hold. Thus, finding a violation of decoding separability with any decoder reflects either a violation of encoding separability or a statistical error. Thus, one strategy that might prove useful in the future is performing several decoding separability tests, each using a different decoder. However, such a test should fulfill two requirements: (1) the decoders should be chosen based on previous work showing that they can detect different violations of encoding separability, and (2) a correction for multiple tests should be applied, to control for the family-wise type I error.

Similarly, we linked encoding separability violations to a pattern difference invariance test by assuming a linear measurement model. This was helpful to prove that in general a violation of encoding separability may or may not lead to a violation of pattern difference invariance, but it is not clear whether some specific measurement models yield a different result. Importantly, it is unlikely that the true measurement model linking neural activity and neuroimaging measurements is simply linear. Thus, more work is necessary to reach a better understanding of exactly what violations of encoding separability can and cannot be detected using the pattern difference invariance test.

### What about encoding modeling?

Faced with the problem of operationalism in the study of neural independence, here our approach has been to propose a very general theoretical framework in which most operational tests can be interpreted and related to properties of encoding such as separability. A different approach would involve building, fitting and selecting among competing encoding models [27]. More specifically, this approach requires building several different encoding models, choosing in each case features such as the number of channels, the shape of the tuning functions, the distribution of neural noise, etc. Encoding separability would hold for some of these models and it would fail for others. The output of each model in terms of neural activities must be linked to neuroimaging data (e.g., estimates of activity from fMRI or EEG) through a formal measurement model. This formal link would allow to derive or numerically approximate a probability distribution of the data given a particular model and parameter set. Once data are obtained, this probability distribution can be used to estimate the parameters of the encoding model and the measurement model that provide the best fit to the data, through maximum likelihood estimation (or Bayesian inference, if priors are added). After estimation, standard model selection procedures (e.g., by AIC, BIC or predictive accuracy in a crossvalidation scheme) would allow to determine what model provides the best explanation for the data. The properties of the chosen encoding model, including its status regarding encoding separability, provide the best explanation for the data.

This approach allows to explicitly test specific features of encoding, and some researchers argue that encoding modeling is the best way to reach valid conclusions about representation in a given brain region from neuroimaging studies [80, 81]. Here we have shown that valid inferences about representation can be made from decoding studies, but we do believe that answering specific questions about representation may be easier through encoding modeling.

However, there are three important challenges faced by anybody wanting to apply encoding models in this way. The first challenge is that this approach will pick the best model among those tested. Thus, a poor selection of competing models will lead to the wrong inferences. Building and fitting encoding models in this way requires an important level of knowledge about what stimulus properties are encoded and how they are encoded [80]. Relatedly, fitting specific models may allow to draw more precise inferences regarding encoding separability, but such inferences cannot be generalized to situations outside those included in the tested models. In contrast, a failure of decoding separability signals a failure of encoding separability regardless of the unknown specific details of the encoding and measurement models. The second challenge is that the process of fitting the models itself may require considerable technical expertise and computational resources. Likelihood functions must be derived or numerically approximated for each model, problems of model mimicry and identifiability must be assessed and solved, simplifying assumptions and constraints must sometimes be placed on the models. The consequences of decisions regarding each of these issues-and the way in which they affect inferences-might not be clear for researchers that are not experienced with modeling. The third challenge has to do with inference and interpretation. It is not always very clear what can and cannot be concluded from the fit of encoding models to data, and recent work suggests that common interpretations of the results of encoding modeling are incorrect [82, 83]. This is complicated further by the fact that many researchers using encoding models are not explicit about their modeling assumptions. For example, many applications involve using multiple linear regression with least squares estimation of weight parameters, where the independent variables are stimulus features assumed to be encoded and the dependent variable is the measured activity in an fMRI voxel or EEG channel. Although never explicitly stated, the assumption behind these models is that there is no neural noise (independent variables in linear regression are not random), the measurement model is linear, and measurement noise is Gaussian and additive. Any conclusion reached using these popular models must be qualified by this set of assumptions.

Overall, we believe that encoding modeling is an excellent way to study the properties of neural encoding using neuroimaging. However, for the reasons outlined before, it seems unlikely to be adopted by experimentalists without a computational background. Indeed, researchers without such a background are probably wise to keep away from it. On the other hand, we have shown here that decoding approaches can lead to valid inferences about the independence of neural representations without being difficult to apply and interpret. We believe that using a decoding separability test offers an improvement over the operational tests of independence commonly used in the literature, without requiring a high level of expertise from researchers.

### Limitations and future work

Our application to face perception research is useful to highlight the kinds of questions that can be answered with the new framework and the type of analysis that should be performed to answer such questions. However, there are limitations of the present study that should be noted. First, results were obtained using a small set of naturalistic stimuli, so they should not be over-generalized. There is no guarantee that the same results will hold for other stimulus sets, and more research is needed before reaching any general conclusion about the separability of identity and emotional expression. Second, our experiment and analyses were performed at the group level. This was done to obtain a statistically-powerful test that is sensitive to violations of separability that are consistent across participants. However, the results may not be representative of individual subjects. We expect that the study of encoding separability at the individual level will require obtaining more data from each participant than what was acquired in the present study.

Our theoretical work might also require further refinement. In particular, the decoding separability test can detect when encoding separability is violated, but it cannot detect when encoding separability holds (see Figure 3).

Decoding separability itself is difficult to prove, as the decoding separability test is a null hypothesis test. Other approaches are necessary to prove the null of decoding separability, such as an arbitrarily small confidence interval around no effect. Such confidence intervals could be computed using resampling methods, such as bootstrapping. Providing evidence favoring the null in this way is usually difficult, as obtaining small confidence intervals requires a large amount of data. Furthermore, there is not much to gain from proving the null of decoding separability, because concluding that decoding separability holds does not lead to conclusions about encoding separability (see Figure 3). Thus, when using the decoding separability test (or any of the other operational tests that we have discussed here), researchers should focus only on obtaining evidence of its failure.

For many researchers, concluding that a dimension is encoded in a separable manner in a given brain region might be considered more interesting; still, an important contribution of our work is showing that this is in general not possible through indirect measures of neural activity or psychophysics. Perhaps specific assumptions about the measurement model producing the data will make it possible to establish a more direct link between decoding and encoding separability, but such assumptions need to be clearly spelled out by researchers, and data should be provided to back them up. One way in which it is possible to directly compare the evidence in favor of encoding separability versus the evidence against encoding separability is within the encoding modeling framework described earlier. As indicated earlier, this framework allows to compare two encoding models that are identical in all respects except their assumption of encoding separability. Unlike in null hypothesis testing, there is no problem with selecting the simpler model in which encoding separability holds, as long as it provides a better explanation for the data.

Although our framework provides a link between neural and perceptual forms of separability, some researchers might consider this link rather weak, as we have only shown that a violation of perceptual separability should be reflected in a failure of encoding separability. Although simply indicating that encoding separability must fail is not very informative about exactly how and where it fails, it is important to understand that here precision has been traded-off for generality. That is, perceptual separability allows to conclude that encoding separability fails, *regardless of how the dimension is encoded by the brain or how it is decoded for performance in a task*. There is a long tradition in vision of linking neural encoding and psychophysics, and more precise conclusions can be reached by making stronger assumptions about encoding and decoding. For example, a common assumption in this literature is optimal decoding through maximum likelihood estimation [84, 85, 86]. The addition of an encoding model that is constrained by results from neurophysiology allows one to make inferences about how many neurons contribute to perception from psychophysical thresholds [84], or about changes in neural tuning functions from changes in threshold-versus-noise functions [86]. Similarly, future research could strengthen the link between neural and perceptual forms of independence for specific dimensions, by including known features of the underlying neurobiology in the encoding models and stronger assumptions about decoding (e.g., optimal decoding), an approach that has not been explored yet in the study of independence.

We must also note that the approach of trading-off precision for generality is entirely within the tradition of how GRT has been developed in the past. That is, most initial work in GRT had the goal of establishing general links between operational tests and different definitions of independence [14, 87, 88, 89]. We believe that this groundwork is necessary before developing more powerful applications to specific problems in vision science.

### Conclusion

The notion of independent processing is central to many theories in perceptual and cognitive neuroscience, but its study has lacked the rigor and integration offered by a formal framework, like the one presented here. This framework allows development of theoretically-driven tests of independence of neural representations, which are more clear and rigorous than the operational tests used thus far. The availability of more rigorous definitions and tests to study separability is likely to advance knowledge in a number of areas in visual neuroscience interested in the notions of independence of processing and representation.

## Materials and Methods

### Participants

Twenty-one male and female right-handed students at the University of California Santa Barbara were recruited to participate in this study. Each participant received a monetary compensation at a rate of US$20/hour. This study was approved by the Human Subjects Committee at the University of California, Santa Barbara, and written informed consent was obtained from all participants.

### Experimental Task

The stimuli and task were identical to those used in a previous behavioral study of separability of face identity and expression [13]. Stimuli were four grayscale images of male faces, part of the California Facial Expression database (http://cseweb.ucsd.edu/~gary/CAFE/). Each face showed one of two identities with either a neutral or sad emotional expression. The faces were shown through an elliptical aperture in a homogeneous gray screen; this presentation revealed only inner facial features and hid non-facial information, such as hairstyle and color.

Participants performed an identification task both outside and inside the MRI scanner. Each stimulus was assigned to a specific response key and the participant’s task was to identify the image presented in each trial. Each trial started with the presentation of a white crosshair in the middle of the screen for 200 ms, followed by stimulus presentation for a single frame (i.e., 16.667 ms at a 60 Hz refreshing rate). Stimulus presentation was short to make it identical to that used in our previous behavioral study. After stimulus presentation, participants pressed a response key; 500 ms later, feedback about the correctness of their response was displayed on the screen for 500 ms (“Correct” in green font color or “Incorrect” in red font color). If the participant took longer than 5 s to respond, the words “Too Slow” were presented on the screen and the trial stopped. Feedback was followed by a variable inter-trial interval, obtained by randomly sampling a value from a geometric distribution with parameter 0.5 and truncated at 5, and multiplying that value by 1,530 ms (the TR value, see below). To obtain estimates of stimulus-related activity with other events in the trial (crosshair and response) unmixed, we used a partial trials design in which 25% of the trials included the presentation of the white crosshair not followed by a stimulus. Participants were instructed to randomly choose a response on these partial trials.

Stimulus presentation, feedback and response recording were controlled using MATLAB augmented with the Psychophysics Toolbox (psychtoolbox.org), running on Mackintosh computers. Participants practiced the identification task on personal Mackintosh computers outside the MRI scanner for about 20 mins. During this training, participants responded on a keyboard. During scanning, participants responded using the Lumina Response Pad System (model LU400-Pair), with the same finger-stimulus mapping as during pre-training.

### Functional Imaging

Images were obtained using a 3T Siemens TIM TRIO MRI scanner with a 12-channel head coil at the University of California, Santa Barbara Brain Imaging Center. Cushions were placed around the head to minimize head motion. A T1-weighted high-resolution anatomical scan was acquired using an MPRAGE sequence (TR: 2,300 ms; TE: 2.98 ms; FA: 9°; 160 sagittal slices; 1 × 1 × 1 mm voxel size; FOV: 256 mm). Additional scans included a localizer and a GRE field map, neither of which were used in the analyses presented here.

Functional scans used a T2*-weighted single shot gradient echo, echo-planar sequence sensitive to BOLD contrast (TR: 1,530 ms; TE: 28 ms; FA: 61°; FOV: 192 mm) with generalized auto-calibrating partially parallel acquisitions (GRAPPA). Each volume consisted of 28 slices (interleaved acquisition, 2.5 mm thick with a 0.5 mm gap; 2.5 × 2.5 mm in-plane resolution) acquired at a near-axial orientation, manually adjusted to cover the ventral visual stream and lateral prefrontal cortex. There were a total of four functional runs per participant (with the exception of five participants who completed three functional runs).

The first run was a standard functional localizer for face regions [76]. Neutral faces, emotional faces and non-face objects were each presented in different stimulus blocks, separated by fixation blocks. Sixteen images of the same type were presented within a stimulus block, each with a duration of 500 ms and a 250 ms inter-stimulus-interval. Fixation blocks consisted of the presentation of a black screen with a white fixation cross in the middle. The sequence started with a fixation block, followed by 6 blocks of each image category (18 total), each followed by a fixation block, for a total of 37 blocks. Blocks lasted for 12 seconds, and the whole scan lasted about 7.5 mins. The order of image types (e.g., neutral-emotional-object) was counterbalanced across blocks. To ensure attention to the stimuli, participants were asked to push a button whenever an image was repeated in the sequence. Four of the 15 stimuli in a block were repetitions, randomly positioned in the stimulus sequence.

In all other functional runs, which lasted about 10 mins each, participants performed the identification task described earlier, without feedback. Each of the four images was repeated 25 times, for a total of 100 trials per run. Stimuli were viewed through a mirror mounted on the head coil and a back projection screen.

### Statistical Analyses

#### Anatomical scans

Processing of structural scans was done using FSL (www.fmrib.ox.ac.uk/fsl), and included brain extraction using BET and nonlinear registration to MNI 2mm standard space using FNIRT nonlinear registration. The inverse transformation was obtained to transform volumes from standard space back to subject space. For visualization purposes, some statistical maps were converted from MNI standard space to the PALS-B12 surface-based atlas using CARET v.5.65 (www.nitrc.org/projects/caret/) and selecting the options *average of the mapping to all multi-fiducial cases* and *enclosing voxel algorithm*.

#### Preprocessing of functional scans

Preprocessing of the functional scans was conducted using FEAT (fMRI Expert Analysis Tool) version 6.00, part of FSL (www.fmrib.ox.ac.uk/fsl). Volumes from all three runs of the main identification task were concatenated into a single series using *fslmerge*. Preprocessing included motion correction using MCFLIRT, slice timing correction (via Fourier time-series phase-shifting), BET brain extraction, grand-mean intensity normalization of the entire 4D dataset by a single multiplicative factor, and a high-pass temporal filtering (Gaussian-weighted least-squares straight line fitting, with sigma=50.0s). The data from the functional localizer were spatially smoothed with a Gaussian kernel of FWHM 4.0mm. The data used in the main separability analysis were not spatially smoothed during preprocessing. Each functional scan was registered to the corresponding structural scan using boundary-based registration (BBR) in FLIRT with default parameters.

#### Neural activity estimates

After preprocessing of the functional scans, estimates of single-trial stimulus-related activity were obtained for the faces in the main identification task. We used the iterative FBR (finite BOLD response) method described by Turner et al. [90] to deconvolve the BOLD activity related to each stimulus presentation. This method avoids assumptions about the hemodynamic response function that are inherent to parametric estimation methods and it is more successful than the latter in unmixing the responses to temporally adjacent events in event-related designs [90]. Instead of assuming a particular shape of the hemodynamic response function, the full shape of the BOLD response to a stimulus is estimated through a set of 12 FBR regressors that are ordered in sequence. In the regression matrix, each event is represented by a set of 12 ones, starting at the beginning of the event. The method is called “iterative” because it iterates through each trial to estimate the BOLD activity related to the stimulus presentation in that trial only. To do this, a group of 12 regressors is created for the target stimulus, and separate groups of 12 regressors are created for the four stimulus classes in the experiment (the target trial was excluded from the regressor of its class), and for the conjunction of crosshair presentation and response. This results in the estimation of 12 regression coefficients for the target stimulus, which are kept while all other regressors are discarded (they are included only to unmix their influence from the target estimates of the BOLD response). As indicated above, the process is iterated for each trial, resulting in a set of spatiotemporal maps (one for each stimulus presentation), representing estimates of the BOLD activity in each voxel and each of 12 TRs starting at the time of stimulus presentation. The algorithm was implemented in MATLAB (The MathWorks, Natick, MA, USA).

#### Decoding separability test

Here we describe the procedures used to implement a decoding separability test. Theoretical results linking this test to the notions of decoding separability, encoding separability and perceptual separability, as well as a justification for the application of this test to neuroimaging data, can be found in the Results section.

Figure 8 is a schematic representation of the decoding separability test. In this simplified example, we consider only two voxels. The estimates of activity are thus represented in a two-dimensional voxel space. Each point represents activity on a different trial, with each color representing a different stimulus that has been repeatedly presented during the experiment. Decoding facial expression from these two voxels using a linear classifier involves finding a hyperplane in the activity space that best separates trials in which a neutral face was shown from trials in which a sad face was shown (the dotted line in Figure 8). The line orthogonal to the classification bound (sometimes called the classifier’s “decision variable”) represents the direction in voxel space that best discriminates one expression from the other. Thus, it is reasonable to assume that this is the direction in this specific voxel space along which expression is encoded. Using that specific direction in voxel space is not a requirement of the decoding separability test to be valid (the only requirement is that a linear decoding scheme is used; see Results section), but it allows us to link the present work to more traditional MVPA techniques. If we take all the observed data points and project them onto this “expression” dimension, we can use these projected points to estimate a distribution of decoded values. *Decoding separability* holds if this distribution of decoded values is invariant across changes in the stimulus on a second, irrelevant dimension. To test for decoding separability, the two distributions of points for a given expression (e.g., the orange and green distributions for “sad”), each corresponding to a different identity, can be compared to one another.

**Figure 8:**
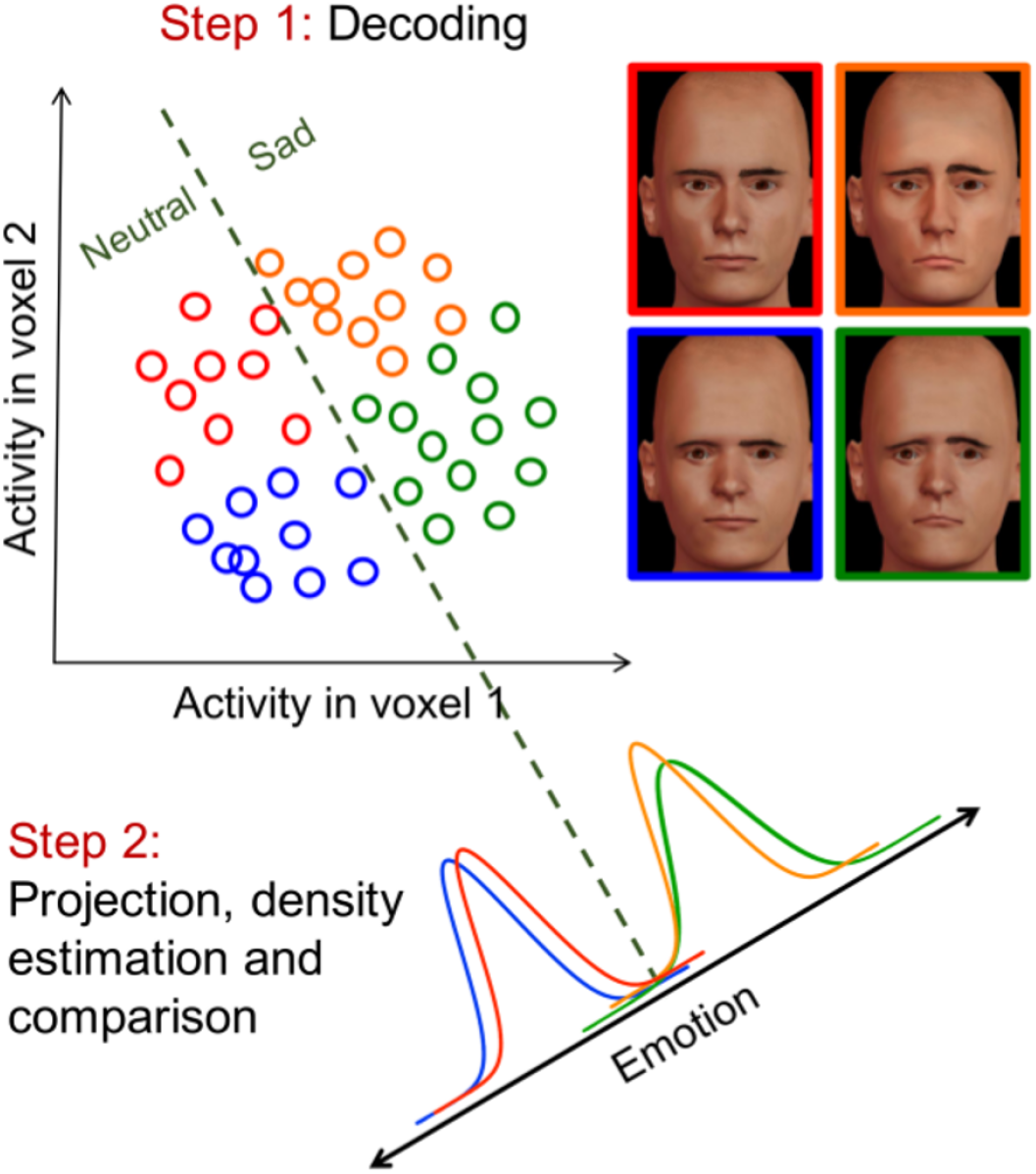
A schematic representation of a test of decoding separability for neuroimaging data, implemented as an extension to traditional linear decoding procedures. The simplified example considers the representation of four stimuli in two voxels. Each point represents activity on a different trial, and each color represents a different stimulus that has been repeatedly presented during the experiment. The dotted line represents a classification bound that separates trials according to emotional expression. The line orthogonal to this bound represents the direction in voxel space that best discriminates one expression from the other. Decoding separability holds if the distributions along this dimension for a given value of the target dimension (emotional expression) are equivalent across changes in the irrelevant dimension (identity). Adapted from: http://figshare.com/articles/Test-of-separabilityof-neural-representations/1385406

In the Results section, we link this decoding separability test to our main theory, and show that it is a valid test of violations of separability in neural representations, even when it is applied to indirect and noisy measures of brain activity, like those obtained from fMRI.

In two separate analyses, we tested the separability of emotion from identity and the separability of identity from emotion; because both analyses are identical, we will describe the analysis in terms of decoding separability of a “target” dimension from an “irrelevant” dimension. Each of the regression coefficients obtained in the previous step were standardized by column. We used a searchlight procedure [77] with a spherical mask that had a radius of three voxels. In each step of the analysis, the searchlight was centered on a different brain voxel and the selected data were used to train a linear support vector machine (SVM, using the *LinearNuSVMC* classifier included in pyMVPA) to decode the target dimension using all the available data. Then the data were projected to the normal line to the classification hyperplane to obtain a number of decoded values on the target dimension. Using Python augmented with the SciPy ecosystem, the group of decoded values for each stimulus was used to obtain kernel density estimates (KDEs) of their underlying probability distribution. A gaussian kernel and automatic bandwidth determination were used as implemented in the SciPy function *gaussianJkde*. Let 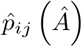 represent the KDE for a stimulus with value *i* on the target dimension and value *j* on the irrelevant dimension, evaluated at point *ς*. The index *i* can take one of two values representing, for example, “sad” and “neutral” when the target dimension is emotional expression, as in the example given in Figure 8. Similarly, the index *j* can take one of two values representing “identity 1” and “identity 2”, as in this example identity is the irrelevant dimension. Then an index of deviations from decoding separability (*DDS*) was computed from all four KDEs obtained from the target dimension (in this example, emotional expression), according to the following equation:

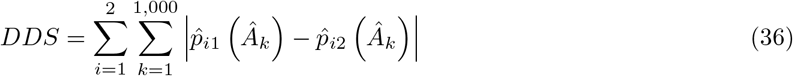

Each KDE was evaluated at 1,000 evenly-spaced points *Â_k_*, indexed by *k* = 1, 2…1,000, starting at the minimum data point minus half the data range, and finishing at the maximum data point plus half the data range. Note that the value 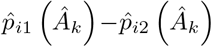 represents the difference between two distributions of decoded values, both related to stimuli with the same value in the target dimension (e.g., “sad”, represented by the index *i*) but different values in the irrelevant dimension (e.g., “identity 1” and “identity 2”, represented by the indexes 1 and 2 in the equation). The index uses the absolute value of the difference between a pair of distributions, which by definition is the *L*1 distance between the two (discretized) distributions (see Equation 27 below). If separability holds, then the distance between distributions should be zero. However, this is only true if we had access to the true distributions. Any error in the KDEs should produce differences between distributions that are added to the *DDS*. This makes it difficult to statistically test for deviations of separability, as the data from multiple participants cannot be combined (differences in the estimation error of the KDEs produces differences in scale of the statistic) and the expected value of the statistic under the null hypothesis is unknown. Under the assumption of decoding separability, two distributions that share the same level of the target dimension but different levels of the irrelevant dimension are identical. That is, data points from those distributions are exchangeable. Taking this into account, we standardized the statistic in the following way: (1) we shuffled the level of the irrelevant dimension for each data point 200 times (separately for each level of the target dimension); (2) each time we computed the *DDS*, yielding an empirical distribution function (EDF) of the statistic under the assumption of decoding separability; (3) the final standardized value was the percentile of the observed *DDS* in the EDF minus 50, representing percentile deviation from the median of the EDF.

Repeating this process for all searchlights resulted in a *DDS* map for each participant, which were converted to the participant’s anatomical space using FSL’s FLIRT linear registration and then to MNI 2mm standard space using FSL’s FNIRT nonlinear registration. The resulting *DDS* maps in standard space were submitted to a nonparametric permutation test using FSL’s *randomise* program [91], with the option *clusterm* for correction for multiple comparisons (which uses the distribution of the maximum cluster mass in the permutation test), a cluster threshold of 2.53 (corresponding to *p*=0.01, uncorrected), variance smoothing with a sigma of 5 mm, and 5,000 permutations.

For visualization purposes, the volumes with significant statistics obtained from the permutation test were converted to the PALS-B12 surface-based atlas using CARET v.5.65 (www.nitrc.org/projects/caret/), and displayed together with the borders of face-selective areas from the localizer scan.

#### Face-selective regions

Face-selective regions were defined using the data from the functional localizer. Low-level analyses were performed separately on the data from each participant. Three explanatory variables (EVs) were defined: Neutral Faces, Emotional Faces and Objects, each corresponding to a boxcar function covering the corresponding blocks in the functional scan (see Functional Localizer description above). These boxcar functions were convolved with the default Gamma hemodynamic response function in FSL, which has a mean lag of 6s and a standard deviation of 3s. A temporal derivative and temporal filtering were added to the design matrix. Two contrasts were formed: Faces (Neutral Faces + Emotional Faces) > Objects, to define regions selective to face information in general, and Emotional Faces > Neutral Faces, to define regions selective to face emotional expression more specifically. Each of these contrasts resulted on a separate map of *z* statistics for each participant. The individual *z* statistical maps were used as input to a high-level analysis, using a mixed-effects model (the option FLAME 1+2 in FSL), to generate a group map for each contrast. Clusters were first identified by thresholding the maps at *z*=2.3; the experiment-wise false positive rate (*α* = 0.05) was controlled by using a threshold on cluster size derived from Gaussian random field theory.

The volumes with significant clusters obtained from the two contrasts were converted to the PALS-B12 surface-based atlas and their borders were manually drawn using CARET v.5.65 (http://www.nitrc.org/projects/caret/). These region borders were used as rough landmarks for the interpretation of the main results of the decoding separability analysis.

The Faces > Objects contrast was additionally used to define face-selective functional regions of interest (ROIs) using the Group-Constrained Subject-Specific (GSS) described by [92]. First, individual maps were thresholded at *p* < 0.05, uncorrected, and the resulting thresholded images were binarized. It was necessary to use a much more liberal threshold than that used by Julian et al. (*p* < 0.0001) to obtain ROI masks in most participants (even at this low threshold, we did not obtain an ROI for the OFA in one participant), because our study was designed to carry out analyses at the group level (see below) and therefore the contrast had less power than that of Julian et al. At the level of individual participants. Second, we took the group-level “parcels” provided by Julian et al. in MNI 2mm standard space (available at http://web.mit.edu/bcs/nklab/GSS.shtml), and transformed them to the participant’s functional space using FNIRT. Third, we intersected the individual binary maps and the group-level parcels to define ROIs corresponding to the fusiform face area (FFA), occipital face area (OFA) and superior temporal sulcus face area (STS) in each individual participant.

Additional anatomical ROIs were obtained, to serve as controls and explore the behavior of our decoding separability test in different conditions. We obtained an ROI corresponding to primary visual cortex (PVC) from the Juelich Histological Atlas, and an ROI corresponding to the lateral ventricles from the Harvard-Oxfor Subcortical Structural Atlas; both atlas are included with FSL. The obtained ROIs were thresholded at a value of 20 using *fslmaths* and binarized. These final ROI masks, which were in MNI 2mm standard space, were transformed to each participant’s functional space using FNIRT.

All ROIs were obtained for both the left and right hemispheres.

## Acknowledgements

This work was supported in part by grant no. W911NF-07-1-0072 from the U.S. Army Research Office through the Institute for Collaborative Biotechnologies, and by NIH grant 2R01MH063760.

